# A multiparametric anti-aging CRISPR screen uncovers a role for BAF in protein translation

**DOI:** 10.1101/2022.10.07.509469

**Authors:** Sophia Y. Breusegem, Jack Houghton, Raquel Romero-Bueno, Adrián Fragoso-Luna, Katherine A. Kentistou, Ken K. Ong, Anne F. J. Janssen, Nicholas A. Bright, Christian G. Riedel, John R. B. Perry, Peter Askjaer, Delphine Larrieu

## Abstract

Progeria syndromes are very rare, incurable premature aging conditions recapitulating most aging features. Here, we report the first whole genome, multiparametric CRISPR anti-aging screen, identifying 43 new genes that can reverse multiple aging phenotypes in progeria. The screen was implemented in fibroblasts from Néstor-Guillermo Progeria Syndrome (NGPS) patients, carrying a homozygous p.Ala12Thr mutation in barrier-to-autointegration factor (BAF A12T). The hits were enriched for genes involved in protein translation, protein and RNA transport and osteoclast formation. We further confirmed that BAF A12T drives increased protein translation and translational errors that could directly contribute to premature aging in patients. This work has highlighted the power of multiparametric whole genome synthetic rescue screens to identify new anti-aging genes and uncover novel biology behind progeria-associated cellular dysfunction.

**One-Sentence Summary:** A whole genome multiparametric screen in progeria identifies new pathways that can reverse cellular aging phenotypes.

## Main Text

Premature aging syndromes (progerias) are very rare conditions that recapitulate many aspects of physiological aging, causing symptoms such as alopecia, lipodystrophy, cardiovascular dysfunction, and bone dysfunction well before the expected age of onset. Many progeria syndromes are caused by mutations in nuclear envelope (NE) associated proteins. For example, the classic Hutchinson-Gilford progeria syndrome (HGPS) is caused by mutations in *LMNA* encoding for the nuclear lamina proteins lamin A and C (*1, 2*). A more recently described progeria, Néstor-Guillermo progeria syndrome (NGPS), is caused by a recessive alanine to threonine amino acid substitution at position 12 (p.Ala12Thr) in the *BANF1* gene, encoding the 10 KDa protein barrier-to-autointegration factor (BAF) (*3, 4*). BAF forms dimers that bind to DNA (*5, 6*), lamin proteins as well as LEM-domain containing proteins of the inner nuclear membrane including emerin (*7, 8*). Through these interactions, BAF exerts critical functions including reformation of the NE after mitosis (*9, 10*) and NE rupture repair (*11*). NGPS patients suffer from multiple aging-associated pathologies, such as alopecia and lipodystrophy, as well as osteoarthritis and joint stiffness. However, the most severe phenotype affecting their quality of life is bone dysfunction, with osteoporosis and dramatic skeletal deformation (*3, 4*). NGPS patients have an increased life expectancy compared to HGPS, probably due to the absence of cardiovascular dysfunction, the main cause of death in HGPS (*12, 13*).

NGPS remains poorly characterized compared to HGPS, and insights into the molecular mechanisms behind the disease have only recently started to emerge (*14–16*). Therefore, with the aim of identifying new genes and pathways relevant to NGPS, as well as potential targets for therapeutic avenues in the disease, we carried out a whole genome CRISPR/Cas9 arrayed microscopy screen in patient derived NGPS cells. We assessed the impact of deleting each of the ∼20,000 human genes on four specific cellular NGPS aging phenotypes simultaneously, to identify “rescue” genes acting across several phenotypes. Through this approach, we identified 43 genes enriched in pathways including protein translation and bone cell development, that improve NGPS cellular phenotypes.

### NGPS fibroblasts show distinct phenotypes compared to HGPS

Fibroblasts from HGPS patients have been extensively studied and characterized. The best-established phenotypes in these cells include NE blebs and invaginations, loss of heterochromatin, downregulation of lamin B1, nucleocytoplasmic transport defects and accumulation of DNA damage foci (*17–22*). To establish whether NGPS cells might share similar phenotypes, we obtained immortalized fibroblasts from two of the three so far identified NGPS patients (NGPS1 and NGPS2) and age-matched immortalized fibroblasts (wild-type – WT) (gift from Dr. Carlos Lopez-Otin). We observed severe NE abnormalities by electron microscopy, including deep folds and NE blebbing but no apparent loss of heterochromatin at the nuclear periphery (Fig. 1A). On the contrary to primary NGPS cells being unable to proliferate in culture (personal communication from Dr. Carlos Lopez-Otin), the immortalized NGPS cells grew at an only slightly lower rate compared to WT cells (Fig. 1B). As previously reported (*4, 23*), we confirmed the accumulation of the NE protein emerin into the cytoplasm of both NGPS1 and 2 (Fig. 1C), and the absence of lamin B1 downregulation (Fig. 1C) and of DNA double strand breaks accumulation (53BP1 foci Fig. 1D, E), both of which being common hallmarks of HGPS cells. In addition, and unlike HGPS cells, NGPS fibroblasts did not display loss of nuclear Ran – used as a reporter for nucleocytoplasmic transport (Fig. 1D, F) or of the heterochromatin markers HP1γ and H3K9me3 (Fig. 1G-I and Fig. S1A-D). We did however observe an increase in the expression of the cyclin-dependent kinase inhibitor p21 – involved in cell cycle progression, apoptosis and DNA damage response – in both NGPS cell lines but to different extents (Fig. 1I and Fig. S1D, E).

**Fig. 1.**
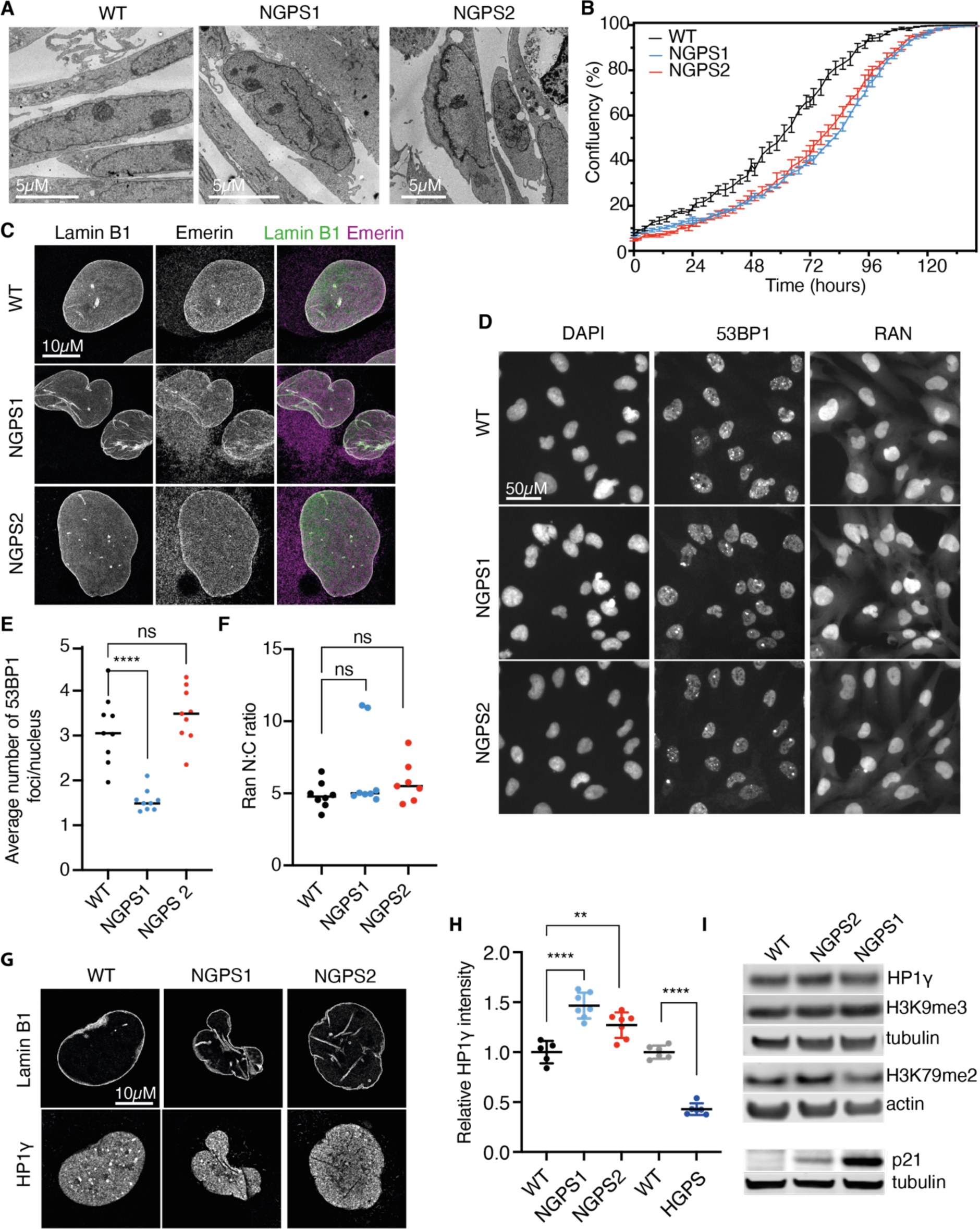
NGPS fibroblasts show distinctive phenotypes to HGPS. (**A**) Representative transmission electron micrographs of WT, NGPS1 and NGPS2 fibroblasts. (**B**) Representative proliferation curves of the indicated cell lines, obtained and analyzed with an Incucyte S3 live-cell analysis system. (**C**) Representative high resolution microscopy images showing the expression and localization of lamin B1 and emerin in WT and NGPS fibroblasts. (**D**) Immunofluorescence images showing DNA damage foci (53BP1) and nucleocytoplasmic transport (Ran) in WT and NGPS fibroblasts. Images were obtained with the CX7 high-content microscope and quantified in (**E**) and (**F**) using the HCS Studio^TM^ software (Data from 2-3 technical replicates, obtained from 3 independent experiments; statistical comparison using unpaired t test). (**G**) High resolution microscopy images of the heterochromatin marker HP1γ in WT and NGPS cells. (**H**) Nuclear HP1γ levels quantified using the high-content microscope in 2 independent experiments, each averaging 500 nuclei in 3 wells. Means ± SD are shown, and unpaired t-tests are used for comparisons. (**I**) Representative western blotting from 3 independent experiments, showing the indicated heterochromatin marks (top blot) and p21 (bottom blot) in NGPS cells compared to WT.

### A combination of four specific NGPS phenotypes amenable to high-throughput screening

Phenotypes associated with NE dysfunction in progeria cells can only be assessed by microscopy, precluding a whole genome pooled CRISPR screening approach. We therefore designed a multiparametric, microscopy based CRISPR screen, aimed at interrogating the whole genome to identify genes that when deleted, could rescue multiple NGPS cellular phenotypes. This approach is based on the principle of synthetic rescue, relying on a genetic interaction whereby a protein not involved in the aetiology of the disease is targeted, thereby rebalancing the pathophysiological state towards a “healthy” one. Our previous work, based on small scale screening, has established the proof-of-concept of this approach in HGPS, where we identified N-acetyltransferase 10 as a new target to reverse HGPS aging phenotypes (*24–26*). The design of the screen is depicted in Fig. 2A and involves multiple steps. The first major step relied on the identification of a combination of phenotypes, specific to the BAF A12T mutation, significantly different in NGPS cells compared to WT cells, and amenable to high throughput screening. To this aim, we took advantage of our recently established NGPS2 isogenic cell lines, in which we reversed the homozygous BAF A12T mutation using CRISPR/Cas9 (NGPS2 corrected) (*14*). The screen set-up also required the design of specific pipelines to reliably identify and quantify these phenotypes. The first of these phenotypes is the delocalization of emerin from the nucleus into the cytoplasm (Fig. 2B-C and Fig. S2A), as characterized by previous studies (*4, 23*). The second phenotype is the increased nuclear deformation, commonly observed in progeria cells (*17, 27*) and in cells from normal aged individuals. This was measured using a perimeter to area ratio (P2A) (Fig. 2D and Fig. S2B). Additionally, we identified an increased number of NE ruptures during interphase in NGPS cells (Fig. 2E and Fig. S2C), further characterized in our recent study (*14*). NE ruptures are identified on fixed cells by the presence of nuclear blebs (observed using LAP2 staining), the first step in the NE rupturing process (*28–30*) (Fig. 2E, F, H). Finally, we observed an increased frequency of micronuclei formation in NGPS cells (Fig. 2G, I and Fig. S2D), a known marker of genomic instability. The combination of these four phenotypes was subsequently chosen for the whole genome screen.

**Fig. 2.**
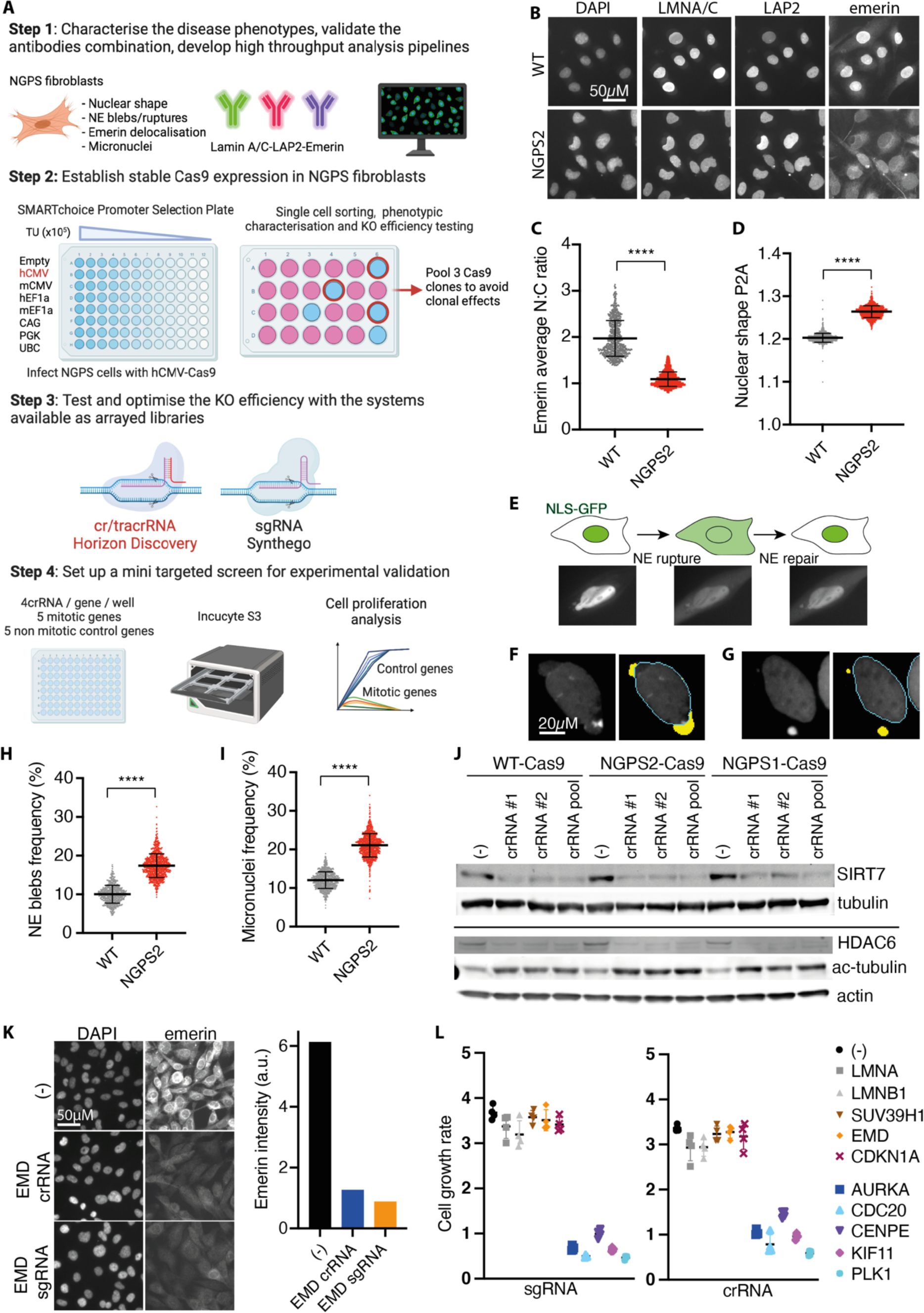
Whole genome CRISPR screening set-up in NGPS fibroblasts. (**A**) Schematic detailing the four main steps involved in the CRISPR screen set-up (created with BioRender.com). (**B**) Example immunofluorescence images of the indicated stainings used in the screen and obtained with the high-content microscope. (**C**) Quantification of the nucleus to cytoplasmic (N:C) emerin intensity ratio in WT and NGPS2 cells. (**D**) Quantification of the nuclear shape using a perimeter to area (P2A) analysis obtained with HCS Studio. Each data point is the average value measured over 500 cells, in 7 experiments, lines indicate the average ± SD and unpaired t-tests were used for statistical analysis. (**E**) Graphical representation and corresponding immunofluorescence images showing the identification of blebs as the origin of NE ruptures using a nuclear localization signal (NLS) reporter tagged with a GFP. (**F**, **G**) Representative nuclei imaged with the CX7 microscope, showing the outlines of the nuclei in blue, and the nuclear blebs (F) or micronuclei (G) in yellow as detected by the HCS software and quantified (as in C and D) in (**H** and **I**). (**J**) Knock down efficiency of individual or pooled crRNAs assessed in the clonal population of Cas9-expressing WT and NGPS cells. (**K**) Representative immunofluorescence images of emerin intensity in NGPS1-Cas9 cells upon transfection of a pool of 3 crRNAs or sgRNAs and quantified using the HCS image studio software (right panel). (**L**) Comparison of cell growth inhibition upon transfection of the indicated sgRNA or crRNA in NGPS1-Cas9 cells. The growth rate was measured using an Incucyte S3 live-cell analysis system in 2 independent experiments. The data shows the representative cell growth from 4 images/well, (average ± SD indicated by lines).

Next, we engineered stable Cas9 expression in WT and NGPS fibroblasts for use in the CRISPR screening. After testing the efficiency of protein transduction using various promoters (Fig. S3A, B) we infected WT and NGPS cells with hCMV-Cas9. Despite a good Cas9 expression level in all three cell lines (Fig. S3C), the knock-out efficiency in individual cells was too heterogenous for screening purposes (Fig. S3D). Therefore, we established individual WT and NGPS Cas9 expressing clones grown from single cells and selected the best clones based on Cas9 efficiency assessed by resistance to 6-Thioguanine after HPRT knock-out (*31*) (Fig. S3E-G). We also assessed Cas9 expression level by western blotting (Fig. S3E) and picked clones combining a high cutting efficiency, and no increase in the level of the DNA double strand break marker γH2AX (Fig. S3E, F). Finally, to avoid obtaining results that could result from a clonal effect, we pooled 3 of the selected clones to generate the WT-Cas9, NGPS1-Cas9 or NGPS2-Cas9 cell populations. The NGPS2-Cas9 was the one used for the screen and was generated by pooling clones D, F and I (Fig. S3E-G). These Cas9 cells showed a good level of protein knock-down upon transient transfection with various crRNAs (Fig. 2J – acetyl-tubulin level serves as an additional readout for HDAC6 knockout efficiency).

Finally, as the project started at a time when the first whole genome CRISPR arrayed libraries were being synthetized, we tested the efficiency of the two non-viral systems available at the time, one relying on a CRISPR/tracrRNA (cr/tracrRNA), the other one on a single guide RNA (sgRNA). After comparing the knock-down efficiency of several genes by western blotting (Fig. S3H) and by immunofluorescence (Fig. 2K and Fig. S3I-L), we set up a mini screen relying on cellular proliferation arrest upon knock-down of five different mitotic genes, compared to 5 non-mitotic genes (Fig. 2L). In both cases, we observed a strong proliferation arrest 3 days after transfection of either the sgRNA (Fig. 2L left panel) or the cr/tracrRNA (Fig. 2L right panel) against the mitotic genes (blue/purple genes), without transfection-associated toxicity (grey/red/yellow genes). Based on cost efficiency, we therefore selected the cr/tracrRNA system for the whole genome screen.

### The multiparametric NGPS screen identifies 43 hits that reverse NGPS phenotypes

We then carried out the primary genome wide screen using the pool of 3 NGPS2-Cas9 expressing clones (NGPS2-Cas9). An overview of the screening approach is depicted in Fig. 3A, and the plate layouts are shown in Fig. S4A-B. The four screening phenotypes - nuclear shape, micronuclei, NE ruptures (blebs) frequency and emerin nuclear intensity - were reduced to two dimensions for each single knock-out and mapped as a Uniform Manifold Approximation and Projection (UMAP). The cluster analysis (Fig. 3B) was used to visualise NGPS2 cells (grey dots) alongside the same parameters measured in control cells (blue dots). The two populations were clearly separated, reflecting the phenotypic difference between the control and the NGPS2 cells. The red dots highlight 112 genes that upon being depleted were identified as “normalising” the combination of NGPS phenotypes. To these, we added 18 genes not yet on the list that appeared in the top 15 of three individual phenotypes (see details in the Methods section). We then carried these 130 genes into a validation screen, in which 3 out of the 4 individual crRNAs used in the primary screen were deconvoluted into individual wells (Fig. 3A). We carried out the validation screen in triplicate and confirmed 43 hits with high confidence (at least 2 out of 3 crRNA and confirmed in the 3 replicates). A gene ontology analysis revealed specific enrichment in biological processes including translation, protein and RNA transport as well as osteoclast development – of high relevance to the patient phenotypes (Fig. 3C, D). In accordance with the concept of synthetic rescue, only 3 hits were proteins associated with the NE: NUP160, SEC13 and AHCTF1 (ELYS).

**Fig. 3.**
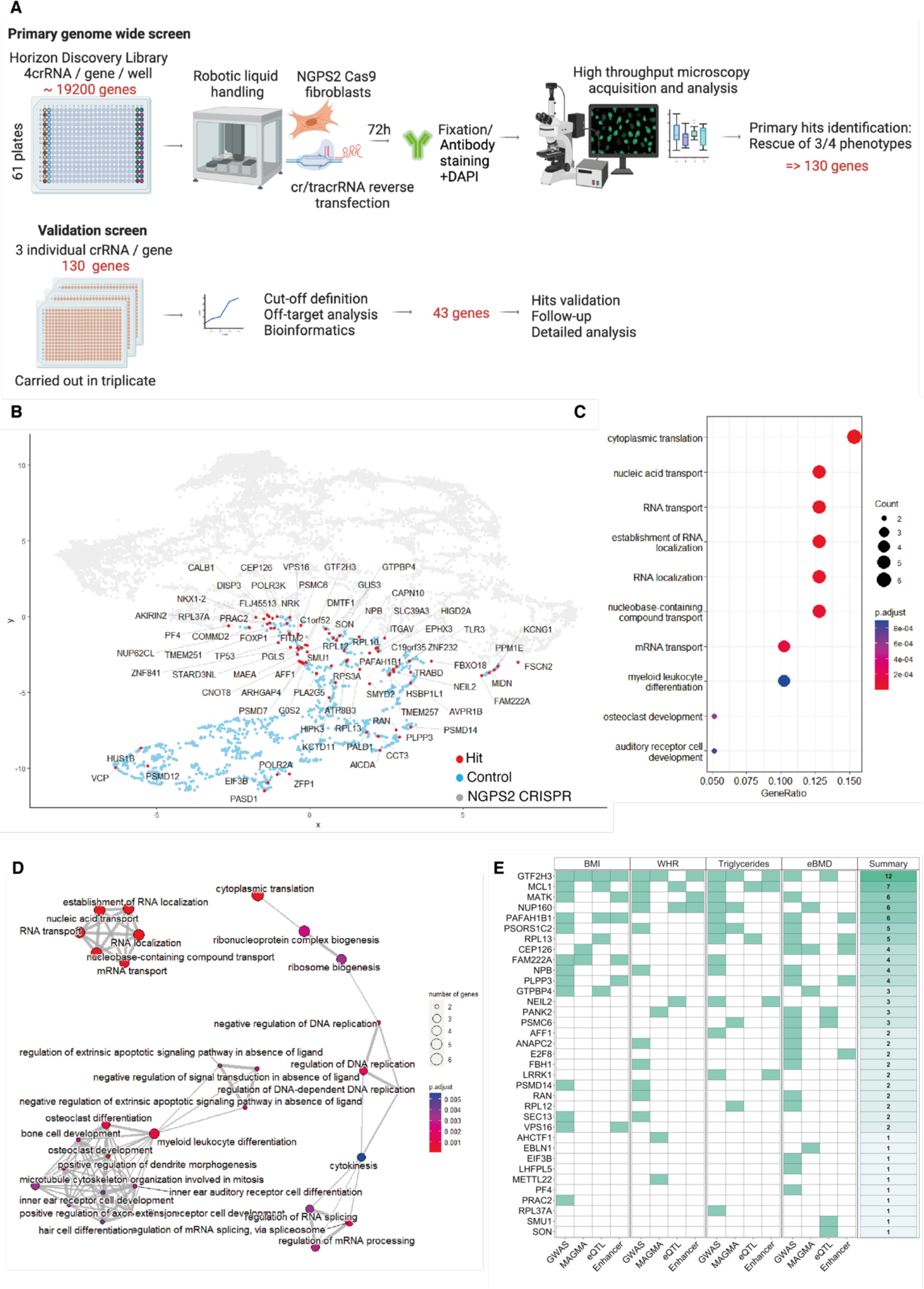
Identification of new genes and pathways normalizing NGPS phenotypes. (**A**) Schematic showing the experimental outline of the primary and validation screens in NGPS2-Cas9 fibroblasts (created with BioRender.com). (**B**) Uniform Manifold Approximation and Projection (UMAP) of the cluster analysis from the primary screen. Nuclear shape, micronuclei and NE blebs frequency quantified in NGPS2-Cas9 cells (grey dots) for each single knock-out were reduced to two dimensions and mapped alongside the same parameters measured in matching control cells (blue dots). The hit genes are labeled in red. (**C-D**) Biological processes gene set enrichment was carried out using the clusterProfiler R package. The 43 validated genes were tested against a full homo sapiens ontology database. (**E**) Heatmap showing the overlap between identified target genes and human genetic datasets. For all four phenotypic traits (BMI (n= 806,834), WHR (n=694,649), triglycerides (n=1,253,275) and eBMD, (n=426,824)), target genes were annotated on the basis of (i) proximity to GWAS signals, (ii) coding- variant gene-level associations to the trait (MAGMA), (iii) colocalization between the GWAS and eQTL data and (iv) the presence of known enhancers within the association peaks. A count of the observed concordant predictors (out of a maximum of 16) is displayed in Summary (right panel). Expanded results can be found in Table S12.

To evaluate the potential relevance of our identified candidate genes to normal variation in related health and disease traits, we interrogated phenotypic and genetic data from available large-scale population studies. These analyses aim to assess the impact of naturally occurring alleles, influencing the function of our candidate genes, on related phenotypes. Since NGPS patients display severe skeletal abnormalities and lipoatrophy, we focused on publicly available Genome Wide Association Studies (GWAS) of body mass index (BMI), waist-hip ratio (WHR) adjusted for BMI, circulating triglyceride levels and estimated bone mineral density (eBMD) in sample sizes up to 1,253,275 individuals (Fig. 3E). We found that 30 of our 43 identified genes were within 500kb of a genome-wide significant signal for at least one of these four traits. Specifically, 13 genes were proximal to BMI signals, 11 to WHR, 19 to eBMD and 13 to triglyceride signals (Table S12 & Fig. 3E). To more directly link these proximal associated genetic variants to the function of our candidate genes, we undertook a number of variants to gene mapping approaches (see methods), including assessment of protein-coding variants and integrating activity-by-contact enhancer maps and expression quantitative trait loci (eQTL) data. These analyses highlighted a number of genes with strong support for a direct involvement in these human phenotypes (Fig. 3E). For example, genetic variants residing within enhancers for *RPL13* and influencing its transcript expression in blood, were associated with eBMD, triglycerides and BMI. Collectively these findings demonstrate that our cellular screens are able to identify genes that influence NPGS-like phenotypes in the normal population.

### Increased protein translation rate caused by BAF A12T is reduced upon depletion of the hits

The screen revealed a strong enrichment for genes involved in protein translation, whose modulation has previously been associated with longevity in various model organisms. To gain further insights into protein translation regulation in NGPS, we first assessed nascent protein synthesis using incorporation of the clickable methionine analogue L-homopropargylglycine (HPG). Upon fluorescent labeling of HPG through a “click” reaction, we observed a significant increase in protein synthesis in NGPS cells compared to both WT cells and NGPS corrected cells (Fig. 4A, Fig. S5A) as well as in wild-type fibroblasts overexpressing BAF A12T compared to BAF WT (Fig. 4B, Fig. S5B). This suggests that the BAF A12T mutation is directly driving increased protein translation, similarly to what has been reported previously in HGPS cells (*33*). In accordance with this data, we observed a modest but significant increase in the nucleolar area of NGPS2 cells compared to NGPS2 corrected cells (Fig. 4C-D), with a similar trend – but without significance – observed in NGPS 1 (Fig. S5C, D). A significant increase was also observed in fibroblasts expressing BAF A12T compared to BAF WT (Fig. 4E). Together with this phenotype, we identified through a dual luciferase mistranslation assay (*34*) that both NGPS1 and NGPS2 cells displayed an increased rate of errors during protein synthesis, as observed by an increased readthrough (Fig. 4F, Fig. S5E).

**Fig. 4.**
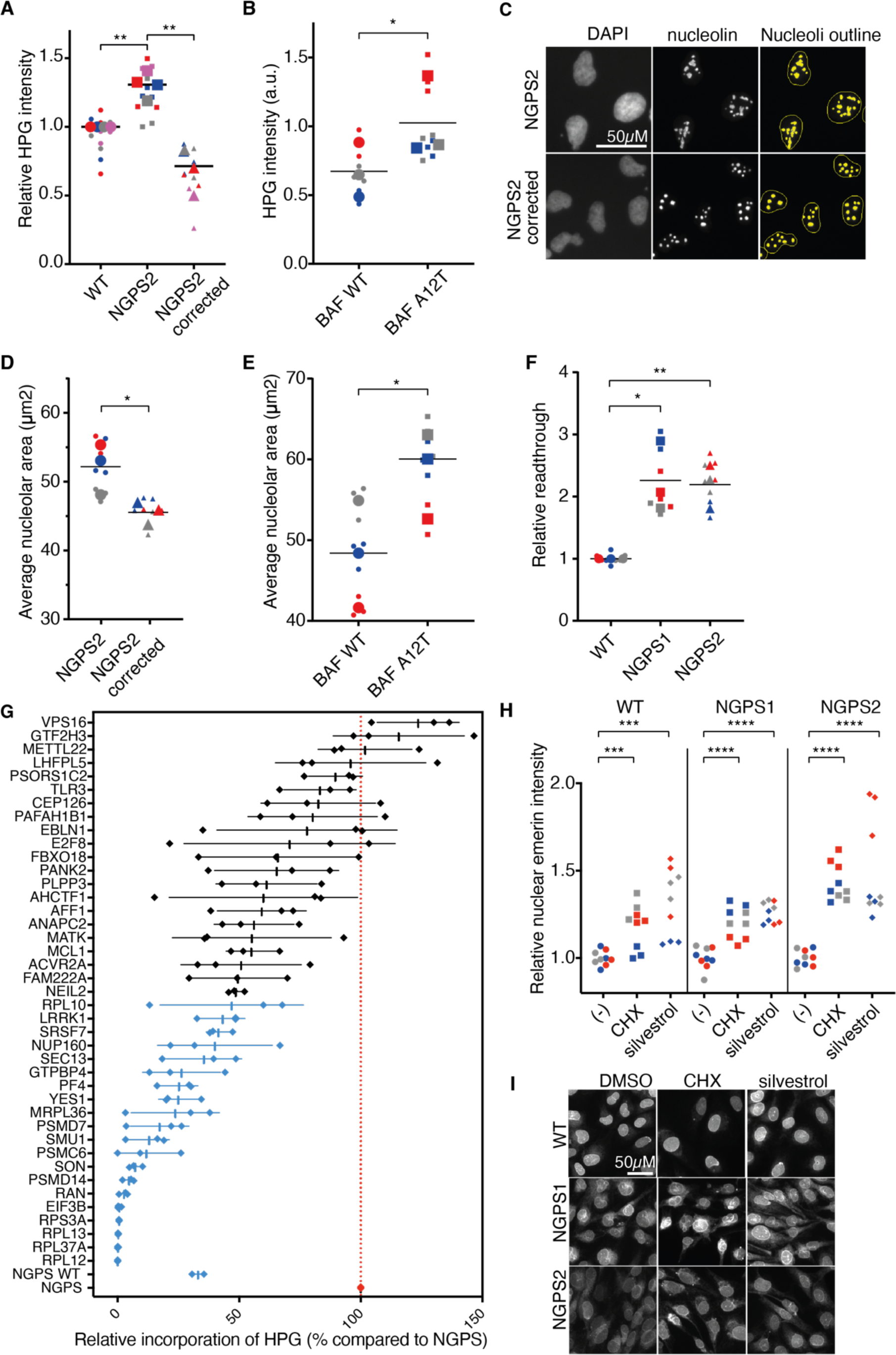
BAF A12T drives enhanced protein synthesis and translation errors. (**A-B**) Nascent protein synthesis assay using HPG incorporation followed by labelling using a “click” reaction with Alexa fluor 488 (AF 488) in WT, NGPS2 or NGPS2 corrected (**A**) or in WT fibroblasts expressing a BAF WT or BAF A12T construct (**B**). AF 488-HPG intensity was quantified using the high-content microscope in 3 independent experiments, each measuring 1000 cells in 3 wells per cell line, graphed as a Superplot with data from different experiments indicated in different colors and larger symbols indicating each experiment’s average. AF 488-HPG intensity was normalized to the corresponding indicated WT cell line, and unpaired t-tests were used for pairwise comparisons between the measured cell lines. (**C**) Immunofluorescence images of nucleolin used to identify and outline the nucleoli (yellow) in the indicated cell lines using the HCS Studio software. (**D-E**) Average nucleolar area quantified in NGPS2 and NGPS2 corrected cells (**D**) or in WT fibroblasts expressing BAF WT or BAF A12T (**E**). Superplots as in (A-B). (**F**) Translation error rate measured as an increased read-through using a dual luciferase assay in 3 independent repeats (Superplot of the data). Results are derived from the ratio hFluc/hRluc, given in fold induction. (**G**) Nascent protein synthesis assay based on fluorescence intensity of HPG AF 488. NGPS2 or NGPS2 corrected cells (NGPS WT) were imaged with the high throughput microscope following siRNA depletion of 41 of the validated screen hits. Depletion of genes highlighted in blue show significant (one-way AOVA with Dunnett’s multiple comparisons test, p<0.05) reduction of HPG incorporation compared to NGPS2 cells transfected with a non-targeting siRNA (red dotted line). Shown are averages ± SD of 3 independent experiments in 500 cells. (**H**) Quantification of emerin nuclear intensity in NGPS1 and NGPS2 cell lines compared to WT, upon protein synthesis inhibition using cycloheximide (CHX) or silvestrol. Three independent repeats are indicated in different colors, and unpaired t-tests are used for pairwise comparisons. (**I**) Representative immunofluorescence images of emerin staining upon protein synthesis inhibition by treatment with cycloheximide (CHX) or silvestrol.

We next asked how depletion of the hits identified in the screen might impact nascent protein synthesis in NGPS cells. To address this question, we used siRNA to deplete 41 of the hit proteins, followed by HPG incorporation and quantification of HGP fluorescent intensity by high-throughput microscopy (Fig. 4G, Fig. S5F). This confirmed a higher rate of protein synthesis in NGPS2 cells (Fig. 4G red dotted line) compared to NGPS2 corrected cells (“NGPS WT”). As expected, depletion of the hits directly involved in protein translation (RPL12, RPL37A, RPL13, RPS3A, EIF3B) almost completely abolished HPG incorporation, and served as a good positive control for the assay. Interestingly however, we observed that depletion of many of the other hits, not known to play a role in protein translation, also led to a reduction of HPG incorporation to various extents in NGPS2 cells, with 21 genes out of the 43 hits (49%) reaching significance (Fig. 4G, blue data points). To assess the potential link between the reduction of protein translation we observed in NGPS2 upon depletion of the hits and the phenotypic improvements identified in our whole genome screen, we treated NGPS1 and 2 cells with the protein synthesis inhibitors cycloheximide (CHX) and silvestrol. Both inhibitors led to the enrichment of emerin into the nucleus of NGPS1 and NGPS2 cell lines (Fig. 4H-I).

### Depletion of RPS3A, PAFAH1B1, VPS16 and SMU1 suppress the lethality of an NGPS ***C. elegans* model**

To establish the potential of our screen hits in translating into improvement of NGPS phenotypes *in vivo*, we used an NGPS *C. elegans* model carrying a *baf-1(G12T)* mutation. The phenotype of these NGPS worms will be described elsewhere, but it is noticeable that control of nucleolar size is impaired as in NGPS cells. When wild type animals were shifted from 16°C to 25°C, nucleolar size was significantly decreased (Fig. 5A-B). Interestingly, this decrease was not observed in *baf-1(G12T)* mutants, which even displayed a mild trend for larger nucleoli at the higher temperature. The *baf-1(G12T)* mutant worms are fully viable, but <10% complete development when combined with an endogenously tagged *gfp::lmn-1*/lamin allele (Fig. 5C, Fig S6A). We took advantage of this sensitized background to test for rescue of developmental arrest upon RNAi-mediated knockdown of 32 genes from our screen (Fig. 5C and Fig. S6B), for which there was a *C. elegans* homologue. Interestingly in regards to the protein translation data we obtained in human cells, depletion of RPS-1 (the *C. elegans* homologue of human RPS3A), a ribosomal protein of the 40S subunit, suppressed the lethality of the *gfp::lmn-1; baf-1(G12T)* animals (Fig. 5B). Three additional genes: *lis-1* (PAFAH1B1) – involved in various dynein and microtubule processes as well as in osteoclast formation (*35*), *vps-16* (VPS16) – a protein involved in protein trafficking to lysosomal compartments (*36*), and *smu- 1* (SMU1) – involved in mRNA splicing (*37, 38*), led to a similar rescue of worm lethality (Fig. 5C). Depletion of the other genes did not rescue the lethality (Fig. S6B), potentially due to some of them – including protein translation genes – being essential *in vivo*.

**Fig. 5.**
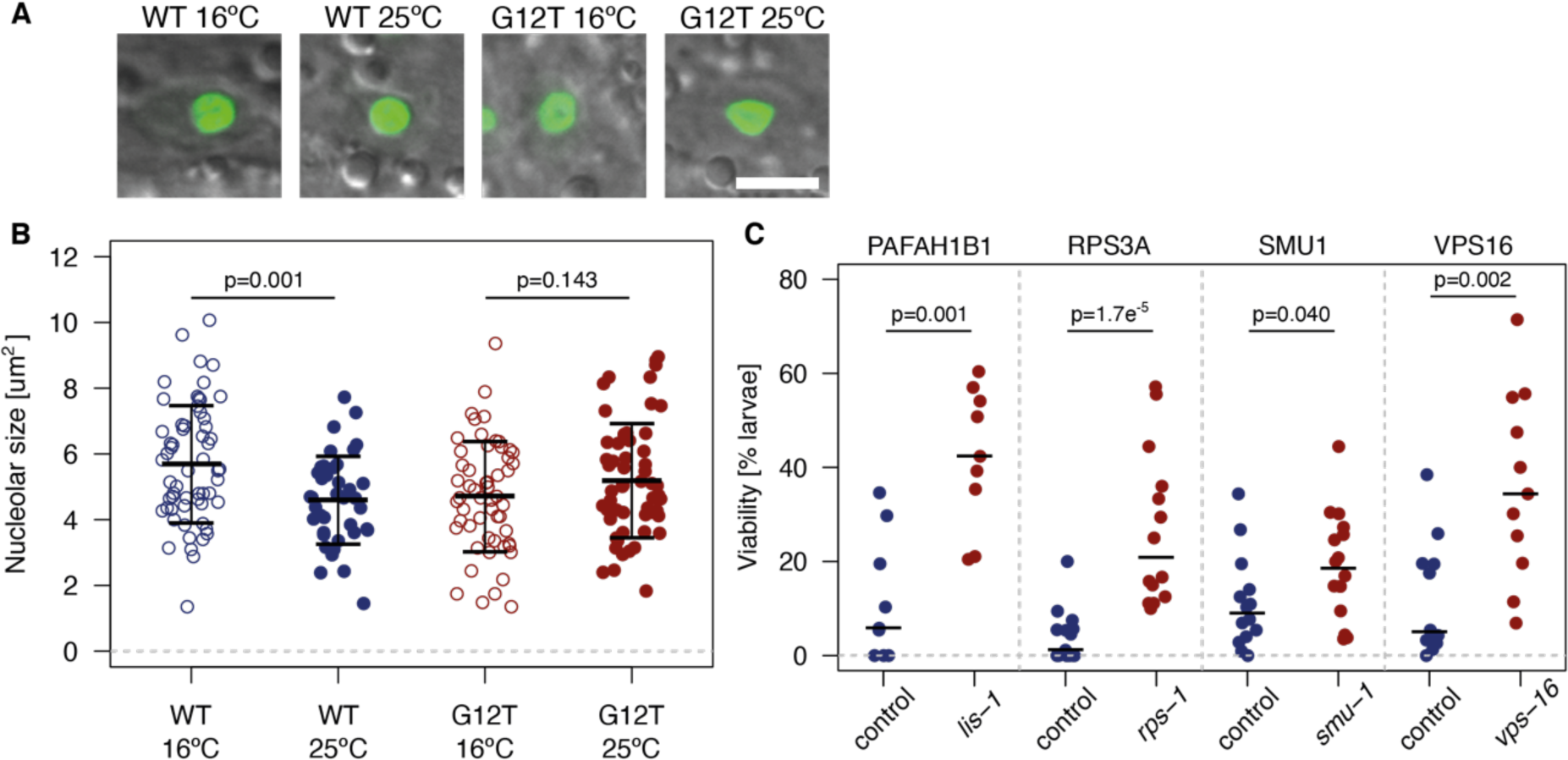
Depletion of PAFAH1B1, RPS3A, SMU1 and VPS16 suppress the larvae lethality of an NGPS *C. elegans* model. (**A**) Confocal images of hypodermal nuclei of live animals expressing FIB- 1::GFP. Shown are overlays of GFP (green) and DIC images. Scale bar 5 um. (**B**) Nucleolar size was measured in wild-type (WT) or NGPS (G12T) *C. elegans* kept at 16 degrees (16°C) or 25 degrees (25°C). The data shows the average nucleolar size ± SD in n=9-11 animals per strain over 2 independent experiments. Unpaired t test p-values were calculated for each genotype. (**C**) The indicated genes were knocked down by RNAi and tested for suppression of lethality in *gfp::lmn-1, baf-1(G12T)* animals. Each point corresponds to the percentage of eggs developing into larvae from a single plate with 50- 100 eggs; 9-15 plates were evaluated in 3-4 independent experiments for each gene. Black lines indicate medians. Mann-Whitney test p-values were calculated for each set of control and test plates.

## Discussion

Our data demonstrates the feasibility and the power of a microscopy based, CRISPR/Cas9 multiparametric synthetic rescue screen for identifying new genes and pathways that can reverse aging phenotypes in progeria cells. While a targeted (320 genes) multiparametric siRNA screen has been carried out previously in a cellular model of HGPS (*39*), our screening approach has interrogated the entire human protein coding genome (19,200 genes) and used NGPS patient cells. By using both WT fibroblasts from an unaffected individual, as well as NGPS fibroblasts in which we reversed the BAF A12T homozygous mutation using CRISPR/Cas9 (*14*), we were able to identify a reliable combination of four phenotypes that were both specific to the BAF A12T mutation, and quantifiable by high throughput microscopy. This approach allowed us to reduce the number of hits to ∼0.2% of the whole genome library, and to identify genes showing the best “rescue” across several phenotypes. In accordance with the principle of synthetic rescue, only 3 of the hits were NE proteins, and more specifically nuclear pore complex proteins: NUP160, SEC13 and AHCTF1 (ELYS). Both SEC13 and ELYS have been previously shown to regulate nuclear size and nuclear import (*40*), a function shared by RAN, another hit from our screen. This suggests that modulating nuclear import might have positive effects on NE organization and function in NGPS cells.

In view of the strong skeletal abnormalities of NGPS patients, it is also worth highlighting that our screen identified two genes involved in osteoclast development: LRRK1 (Leucine-rich repeat serine/threonine-protein kinase 1) and PAFAH1B1/LIS1 (Platelet-activating factor acetylhydrolase IB subunit alpha), whose depletion improve NGPS cellular phenotypes and in the case of LIS1 also increases viability of NGPS worms. LRRK1 has been involved in the regulation of osteoclast activity and bone resorption (*41–43*). Accordingly, LRRK1 dysfunction in mice or human is associated with severe osteopetrosis (*44*). Similarly, PAFAH1B1 regulates osteoclast formation and bone homeostasis through interacting with the protein Plekhm1, involved in osteoclast secretion. PAFAH1B1 depletion has been shown to strongly reduce osteoclast formation (*35*). Therefore, we can hypothesize that the severe osteoporosis and osteolysis observed in NGPS cells might be associated with an increased level or activity of LRRK1 and/or PAFAH1B1. This might contribute to the aberrant NGPS cellular phenotypes that become normalized upon depletion of these genes.

One of the top functional enrichments in the screen was protein translation, with 7 out of the 43 hits being associated with this process. Loss of protein homeostasis is one of the hallmarks of aging (*19*) and inhibition of protein translation has been associated with enhanced longevity in several animal models (*45–47*). Identifying that inhibition of this pathway was beneficial to NGPS cells and NGPS worms, therefore reinforce the current hypothesis that loss of cellular homeostasis observed in both premature and physiological aging might occur through similar mechanisms. More specifically, we observed that the NGPS associated BAF A12T mutation was causing a higher rate of nascent protein synthesis. This could be a common mechanism involved in premature aging, as increased protein synthesis was also observed in HGPS cells (*33*). In the case of NGPS, we saw that this was associated with a higher rate of errors, as seen by an increased stop codon readthrough.

There is currently no mouse model for NGPS but a recent study reported the effect of error-prone protein synthesis in mice (*48*). Through engineering a specific ribosomal mutation, the authors induced genome-wide translational errors in mice. This resulted in reduced lifespan and premature aging features resembling the phenotypes observed in NGPS patients including chest and spine deformation as well as loss of fat. This suggests that deregulation of protein synthesis and accumulation of errors might directly contribute to the premature aging phenotypes of NGPS patients. Therefore, identifying non-toxic pharmacological interventions to slow down protein translation, thereby potentially limiting error-prone translation might yield phenotypic improvement at the level of the individuals. According to our data, this could be achieved through targeting proteins directly involved in protein synthesis (such as RPL proteins) as well as proteins involved in various other cellular functions (Fig 4G). Indeed, in the cellular context of NGPS, we observed a reduction of nascent protein synthesis upon depletion of genes that don’t have any known protein synthesis function. The mechanisms behind this and the link between protein translation and NE integrity remain unknown and will be the basis of future studies, but it does suggest a common mechanism by which depletion of the hits from the screen improve NGPS phenotypes.

Altogether, our screen has shed light on new pathways involved in cellular dysfunction caused by the BAF A12T mutation in NGPS patient cells. This work has also identified potential new therapeutic targets for this incurable premature aging syndrome, supported by the data we obtained *in vivo*, showing rescue of *C. elegans* lethality upon knocking down some of the hits from the screen.

## Acknowledgments

We would like to thank Prof. Lopez Otin for providing us with the NGPS and WT immortalized fibroblasts, Dr Daniel Starr for providing us with the *gfp::lmn-1* alleles, Dr Dimitri Shcherbakov and Dr Erik Böttger for sharing the dual luciferase mistranslation assay plasmids and Dr Dimitri Shcherbakov for helpful discussions on setting up and interpreting the assays, Matthew Gratian (CIMR microscopy) for his help with the high content microscope and screen set up, Henri Huppert (ThermoFisher) for his help with the HCS studio pipelines and Dr Gabriel Balmus for giving us access to the CyBio Felix robot.

## Author contributions

Conceptualization: SYB, DL

Methodology: SYB, JH, AFJJ, AFL, RRB, CR, KK, NB, JBBP, DL

Investigation: SYB, JH, AFJJ, AFL, RRB, KK, NB Visualization: SYB, AFJJ, AFL, RRB, KK, NB, PA, DL

Funding acquisition: DL, PA, JBBP Project administration: DL Supervision: DL, PA, JBBP Writing – original draft: SYB, DL

Writing – review & editing: SYB, JH, AFJJ, KK, JBBP, PA, NAB, DL

### Data and materials availability

All data are available in the main text or the supplementary materials.

## Supplementary Materials

Materials and Methods

Figs. S1 to S6

Tables S1 to S4

Data S1

References (*49–68*)

## Supplementary Materials for

### Materials and Methods

#### Cell culture

Wild type (WT), NGPS5796 (NGPS 1) and NGPS5787 (NGPS 2) immortalized fibroblasts were a kind gift of Prof. Carlos López-Otín (University of Oviedo, Spain). WT fibroblasts were derived from an age-matched healthy individual (AG10803, Coriell repositories) and all three cell lines were immortalized with SV40LT and TERT. Cells were grown at 37°C in a 5% CO_2_ incubator in Dulbecco’s modified Eagle’s medium containing 4.5 g/L glucose (Sigma) supplemented with 2 mM L-glutamine (Sigma), 10% foetal bovine serum, 100 units/mL penicillin and 0.1 g/L streptomycin (complete medium). WT cells expressing Flag-BAF or Flag-A12T BAF were described before (*14*) and maintained in complete medium containing 100 μg/mL hygromycin B (Toku-E, #H007, made up to 100 mg/mL in PBS). Cells were passaged twice a week and used within 12 passages. Cells were free of mycoplasma as assessed using strips (InvivoGen #rep-mys- 20). For the phenotypic comparison with HGPS (Figs. 1 and S1), WT cells were GM05565 fibroblasts and HGPS cells were AG11513 cells, both obtained from Coriell Institute for Medical Research (Camden, New Jersey, U.S.A.).

#### Antibodies and Reagents

Table S1 Lists antibodies and reagents used in Western blotting (WB) and immunofluorescence (IF) applications.

#### Immunofluorescence

Cells were seeded on thickness 1 ½ round coverslips (Thermo Fisher Scientific) in 12-well plates for full-field imaging, thickness 1 ½ square (18 mm × 18 mm) coverslips (Zeiss) in 6-well plates for super-resolution imaging or in a 96-well view-plate (PerkinElmer #6005182) for high-content imaging and grown to a confluency level consistent across the cell lines to be compared. All subsequent steps were carried out at room temperature. Cells were fixed with 4% paraformaldehyde in PBS (Boster, AR1068) for 20 minutes, followed by permeabilization in 0.2% Triton X-100 in PBS for 12 minutes. Unspecific antibody binding was blocked by incubating in 5% BSA (Sigma #A7906), 0.1% Tween-20 in PBS (IF blocking buffer) for 30 minutes. Primary antibodies were diluted in IF blocking buffer (see Table S1) and incubated for 1-2 hours. After 3 washes with PBS cells were incubated with secondary antibodies and DAPI diluted (see Table S1) in IF blocking buffer. After 3 washes with PBS coverslips were mounted using Prolong Gold (P10144) and left to set at room temperature overnight; cells stained in 96-well plates were overlaid with 100 μL/well fresh PBS and stored at 4°C until imaging.

Wide-field immunofluorescence imaging was on an upright Axioimager Z2 (Zeiss) using a 63x 1.4 N.A. oil immersion objective. Super-resolution imaging was on an Elyra PS1 structured illumination microscope (Zeiss) using a 63x 1.4 N.A. oil immersion objective. High-content imaging was on a CellInsight CX7 microscope (Thermo Fisher Scientific) using a 20x 0.35 N.A. objective. HCS Studio software (Thermo Fisher Scientific) was used for quantitative image analysis. For quantitation of H3K9me3 or HP1γ nuclear intensities the DAPI images were used to define the nuclear contour, and a fixed intensity threshold was set to define the stained heterochromatin domains.

#### Western blotting

Confluent monolayers of cells in 6-well plates were washed with PBS and scraped in 70 μL/well SDS lysis buffer (4% SDS, 20% glycerol, and 120 mM Tris-HCl (pH 6.8)). Lysates were incubated for 5 min at 95°C. The DNA was sheared by syringing 10 times through a 25-gauge needle. Absorbance at 280 nm was measured (NanoDrop, Thermo Fisher Scientific) to determine protein concentration. Samples were prepared in NuPAGE sample buffer (Thermo Fisher Scientific #NP0007) and DTT (100 mM) and heated at 95°C for 10 min. Proteins were loaded on NuPAGE 4-12% Bis-Tris gels (Thermo Fisher Scientific), separated in NuPAGE MES SDS running buffer (Thermo Fisher Scientific #NP0002) and transferred to 0.2 μm pore size nitrocellulose membranes (Amersham, #10600004) for immunoblotting. Blotted proteins were reversibly stained with Ponceau S solution (Thermo Fisher Scientific, # A40000279) to allow cutting strips for individual antibody incubations. Membrane strips were first blocked for 30 min in 5% milk in TBST buffer (20 mM Tris, 150 mM NaCl, 0.1% Tween-20) before incubation with primary antibodies diluted in TBST buffer (see Table S1) for 1h at room temperature. After 3 washes with TBST buffer membrane strips were incubated for 1 h with IRDye-conjugated secondary antibodies (LI-COR, see Table S1). After 3 more washes with TBST buffer membrane fluorescence was scanned on an Odyssey CLx imaging system (LI-COR).

#### Transmission electron microscopy

Cell monolayers were fixed by the addition 2.5% glutaraldehyde / 2% paraformaldehyde in 0.1M Na cacodylate buffer, pH 7.2 at 37°C. The monolayer was then scraped from the tissue culture plastic, pelleted in a benchtop microfuge and allowed to cool to room temperature.

The cell pellet was washed with 0.1M Na cacodylate buffer, pH 7.2; post-fixed in 1% osmium tetroxide in 0.1M Na cacodylate buffer, pH 7.2, for 1 hour, and washed with 0.05M Na maleate buffer, pH 5.2. Next, the cells were stained en bloc with 0.5% uranyl acetate in 0.05M Na maleate buffer, pH 5.2, for 1h at 4°C; washed with 0.05M Na maleate buffer, pH 5.2; dehydrated in a graded series of ethanol and exchanged into 1,2-epoxy propane. The cell pellet was then infiltrated with 50:50 epoxy propane: Agar 100 resin overnight before exchange into Agar 100 resin. Finally the cell pellets were embedded in Agar 100 resin in BEEM capsules (Agar Scientific, Stansted, UK) overnight at 60°C.

Ultrathin sections (60nm) were cut using a diamond knife mounted on a Leica Ultracut UC7 ultramicrotome (Leica, Milton Keynes, UK), collected on formvar-coated copper EM grids and stained with uranyl acetate and Reynolds lead citrate. The sections were observed in an FEI Tecnai G2 Spirit BioTWIN transmission electron microscope (Eindhoven, The Netherlands) at an operating voltage of 80 kV. Images were captured using a Gatan US1000 CCD camera.

#### Generation of Cas9-expressing cells

Stable Cas9 expression was engineered in the WT and in both NGPS cell lines as follows. We first assessed protein expression efficiency from seven constitutive promoters (hCMV, mCMV, hEF1α, mEF1α, CAG, PGK and UBC) using a SMARTchoice promoter selection plate (Horizon Discovery, #SP-001000-01). The hCMV promoter showed the highest expression and was chosen to drive Cas9 expression. WT and NGPS cells were transduced with purified lentiviral particles, containing a vector encoding the S. pyogenes Cas9 nuclease under the control of a hCMV promotor according to the manufacturer’s protocol (Horizon Discovery, #VCAS10124). Cells stably expressing Cas9 were selected using 5 μg/mL blasticidin (Merck/Sigma Aldrich), and several clones were isolated from the polyclonal population by single cell sorting. Individual clones were assessed for Cas9 expression, cellular morphology, effect on cell health markers such as DNA damage (γH2AX level) and proliferation, as well as Cas9 cutting efficiency. The latter was assessed based on resistance to 6-thioguanine upon HPRT knock out as described before (*31*) as well as by knock down efficiency using immunofluorescent staining or Western blotting (see CRISPR mini-libraries and Table S2). Based on these, to avoid any clonal effect that might arise from a single clone, three Cas9-expressing clones were selected for each cell line and pooled for further CRISPR experiments (WT-Cas9 or NGPS-Cas9) including the mini-libraries and genome-wide screens.

#### CRISPR mini-libraries

Two CRISPR mini libraries were obtained; one contained pools of 3 sgRNAs targeting 10 genes (Synthego, Redwood City, CA, USA) (Table S3), the other contained pools of 4 crRNAs targeting 20 genes (Horizon Discovery, Waterbeach, UK) (Table S5).

For immunofluorescence or cell proliferation assays, WT-Cas9 or NGPS-Cas9 cells were reverse transfected with the CRISPR reagents in a 96-well view-plate (Perkin Elmer #6005182). Per well 20 μL transfection mix was first prepared in a V-bottom 96-well plate (Greiner #651161) as follows. For crRNA:tracrRNA transfection, 2.5 μL 1 μM crRNA and 2.5 μL 1 μM trRNA were added to 5 μL Optimem; for sgRNA transfection 2.5 μL 1 μM sgRNA was added to 7.5 μL Optimem. To each well containing 10 μL diluted crRNA:tracrRNA or sgRNA was then added 10 μL of a Dharmafect-1 dilution (0.05 μL Dharmafect-1 in 9.95 μL Optimem). The solutions were mixed by pipetting up-and-down and incubated for 20 minutes at room temperature before arraying in a 96-well view-plate (Perkin Elmer #6005182). Meanwhile, cell suspensions were counted using a Countess cell counter (Thermo Fisher Scientific) and dilutions prepared such that to each well were added 3300 cells for WT-Cas9 and NGPS2-Cas9 or 5000 cells for NGPS1 cells in a 80 μL volume in antibiotic-free growth medium. Plates were shaken by hand and placed in a 37°C incubator for 65-72 hrs. For cell survival analysis plates were put in an Incucyte live-cell imaging incubator 24 hrs post-transfection (see *Cell proliferation*).

For Western blotting analysis of sgRNA-transfected cells, reverse transfections were carried out in 12-well format. Per well 3 μL 10 μM sgRNA was diluted in 50 μL Optimem, incubated for 5 minutes before adding diluted Dharmafect-1 (0.6 μL Dharmafect-1 in 50 μL Optimem). Transfection mixes were overlaid with 900 μL antibiotic-free growth medium containing 120,000 NGPS1-Cas9 cells. Plates were shaken and put at 37°C for 72 hrs before cell lysis.

#### Cell proliferation (Incucyte)

WT or NGPS cells were seeded onto 24-well plates (Falcon, # 353047), placed in a live-cell imaging incubator (Incucyte S3, Essen, Germany) and imaged with a 10x phase objective every 4 hours for 96 hours. Incucyte S3 analysis software was used to measure and quantify cell density over time. For assaying the mitotic gene knock-outs in the CRISPR mini-libraries WT-Cas9 or NGPS-Cas9 cells were reverse transfected in 96-well view-plates (Perkin Elmer #6005182) and placed in the Incucyte imaging incubator 24 hours post-transfection.

#### CRISPR library

A genome-wide CRISPR library was obtained from Horizon Discovery (GP-004650-E2-01, GP- 004675-O2-01 and GP-006500-O2-01). The library was arrayed in 61 384-well plates and contained 4 unique crRNAs per gene per well, targeting a total of 19,127 human genes (Table S6). Some crRNA pools were present in duplicate or triplicate, providing internal controls for the screen. The 0.1 nmole crRNA pools were resuspended in 20 μL 10 mM Tris buffer (Horizon Discovery #B-006000-100) to yield a 5 μM master plate. This master plate was further aliquoted into mother plates. To one mother plate containing 2.4 μL/well crRNA was added 21.6 μL 0.56 μM tracrRNA (Horizon); the resulting 24 μL of 0.5 μM crRNA:tracrRNA solution was then divided over transfection-ready daughter plates, each containing 6 μL of 0.5 μM crRNA:tracrRNA per well. All liquid-handling steps were carried out using a CyBio Felix liquid handling robot (Analytik Jena, Jena, Germany). Plates were spun for 2 minutes at 300 x g, sealed using clear polypropylene seals (Starlabs E2796-0793) and stored at -30°C.

#### Genome-wide CRISPR screen

The genome-wide CRISPR screen was carried out with the NGPS2-Cas9 pool of 3 clones. Library plates containing transfection-ready crRNA:tracrRNA complexes were thawed and spun at 300 x g for 3 minutes. Emerin and non-targeting crRNA:tracrRNA complexes were added to columns 23 and 24 (see plate layout, **Fig. S3A**) as positive controls for the transfection efficiency and negative controls respectively. For reverse transfection, Dharmafect-1 (Horizon Discovery) was diluted in Optimem (Thermo Fisher) (120 μL in 23.88 mL Optimem) and incubated for 5 minutes at room temperature before distributing in a 384-well plate. A liquid handling robot (CyBio Felix, Analytik Jena, Germany) was used to add the Dharmafect 1 solution to the crRNA:tracrRNA complexes and to mix the 2 solutions. After 20 minutes of incubation at room temperature, the transfection mixes were distributed in 384-well imaging plates (Greiner #781182) and overlaid with 1200 NGPS2-Cas9 cells/well in 30 μL of complete DMEM medium. WT cells and NGPS2 WT cells (from the 2 clones in which the BAF A12T mutation was reversed using CRISPR Cas9 (*14*) – see Fig. S3A) were added in columns 1 and 2 of each plate (that do not contain transfection complexes). Plates were then kept in a 37°C incubator with 5% CO_2_ for 72 hours.

All subsequent steps were carried out at room temperature. Cells were fixed using a Wellmate robot (Thermo Fisher) by adding 30 μL/well of 4% PFA for 30 minutes. Fixative was removed through plate inversion, cells were washed once with 40 μL/well PBS and permeabilized with 40 μL/well of 0.2% Triton X-100 in PBS for 12 minutes. Unspecific antibody binding was blocked by incubation for 30 minutes with IF blocking buffer (see *Immunofluorescence*). Next, cells were incubated for 1 hour with a mixture of primary antibodies (20 μL/well) diluted 1:1000 in IF blocking buffer: rabbit anti-emerin, mouse IgG_2b_ anti-lamin A/C and mouse IgG_1_ anti-LAP2 (details in Table S1). Following a wash with PBS (40 μL/well), cells were incubated for 1 hour with a mixture of secondary antibodies (Alexa Fluor 488 anti-mouse IgG_2b_, Alexa Fluor 568 anti-mouse IgG_1_ and Alexa Fluor 647 anti-rabbit – details in Table S1), as well as DAPI (Table S1), in IF blocking buffer (20 μL/well). After a final wash with 40 μL/well PBS the cells were overlaid with 40 μL/well fresh PBS, plates were sealed with a black seal (Perkin Elmer) and loaded onto a OrbitorRS plate handling robot (Thermo Fisher) for loading into and imaging using a CellInsight CX7 high-content microscope (Thermo Fisher). 4-channel images were acquired with fixed exposure times using a 20x 0.35 N.A. air objective for 9 fields/well or until 500 nuclei (“objects”) were detected in the DAPI acquisition channel. The HCS studio colocalisation bio-application was used to calculate a nuclear shape parameter for the DAPI-defined objects and the emerin intensity in both the nucleus, defined by the DAPI mask, and in a cytoplasmic ring defined by expansion of the DAPI mask. For micronuclei analysis, the LAP2 images were run through the HCS studio micronuclei bio-application. This analysis module was further modified to detect nuclear blebs, again using the LAP2 images (*49*). All primary screen results are collated in Table S7.

#### Primary screen hit identification

Two multivariate analyses were carried out in R using custom scripts. For both analyses, data dimension reduction was achieved using UMAP (*50*) and clustering using *k* means, with optimal cluster number determined by differential grouping of WT control samples. A four-parameter (average nuclear emerin intensity, micronuclei frequency, nuclear shape factor P2A, and nuclear bleb frequency) analysis, including a scaling step prior to dimension reduction (subtraction of the values of each parameter by the mean of that parameter followed by division of the values of each parameter by the standard deviation of each parameter), yielded 51 hits. A three-parameter analysis, excluding emerin nuclear intensity as few gene knockouts rescued the phenotype, yielded 68 hits. Of the combined 119 hits we excluded genes whose knockout negatively affected cell survival or proliferation (nuclear count < 200), and we added genes not yet on the list that appeared in the top 15 of three individual phenotypes (nuclear shape factor, micronuclei or nuclear envelope bleb frequency), bringing the total number of hits taken forward for a deconvoluted validation screen to 130 (Table S8).

#### Validation screen

For each of the 130 primary screen hits, 3 crRNA sequences were arrayed randomly in the central wells of two 384-well plates as a custom library (Horizon Discovery) (Table S9). Control crRNA:tracrRNA complexes were added to the plates as detailed in Fig. S3B. Transfection, fixing, staining and imaging were all carried out as for the primary screen and repeated 3 times. Aggregate z-scores were calculated for the 3 sequences/genes for each phenotype (Table S10). Genes whose z-score was below (for micronuclei frequency, NE bleb frequency and nuclear shape parameter P2A) or above (for emerin nucleus:cytoplasm intensity ratio) the z-score for the NT control in the 3 repeat experiments were considered validated. The 43 validated genes are in the STRING diagram in Fig. S3C.

#### Nascent protein synthesis assay

Nascent protein synthesis was assayed using incorporation of the methionine analogue L-homopropargylglycine (HPG). Cells were grown in methionine-free medium (Thermo Fisher Gibco #21013-024) for 1 hour before incubation with 1 μM click-iT HPG (Thermo Fisher #C10428) in methionine-free medium for 1 hour at 37°C. All subsequent steps were carried out at room temperature. Cells were fixed with 4% paraformaldehyde in PBS for 20 minutes, followed by permeabilization with 0.2% Triton X-100 in PBS for 12 minutes. Next, a click reaction was allowed to proceed for 1 hour by incubation with 4 mM CuSO4, Alexa Fluor 488 azide (1:1000 dilution) and 10 mM sodium ascorbate in the dark. Click reagents were washed off and nuclei were stained with 0.2 μg/mL DAPI before imaging on the high-content microscope.

#### siRNA transfection

A custom siRNA mini-library containing On TargetPLUS siRNA smartpools (Horizon Discovery) targeting 41 of the 43 validated hits as well as some internal control genes (EMD, LMNA and TMPO (encoding LAP2)) arrayed in a 96-well plate was obtained (Table S11, see Fig. S4F for the plate layout). The 0.1 nmole siRNA/well was diluted to 2 μM with 10 mM Tris buffer (Horizon Discovery #B-006000-100). Transfection mixes were prepared by adding diluted Lipofectamine RNAiMAX (Thermo Fisher Scientific) (1 μL in 33 μL Optimem) to diluted siRNA (5 μL 2 μM siRNA added to 28 μL Optimem) in a V-bottomed 96-well plate (Greiner #651161), mixing and incubating for 10 minutes at room temperature. 20 μL of the transfection mix was then transferred to a 96-well viewplate well (Perkin Elmer #6005182) and overlaid with 80 μL cell suspension (NGPS2 cells, 3000 cells/well). siRNA transfection was allowed to proceed for 72 hrs before cells were assayed for nascent protein synthesis as detailed above.

#### Protein translation inhibition studies

WT or NGPS cells were seeded in 96-well imaging plates (PerkinElmer #6005182) and treated for 72 hours with 0.125 μg/mL cycloheximide (CHX, Sigma # C4859) or 2.5 nM silvestrol (Biovision #B2417-100) before staining and imaging as in the screens.

#### Protein mistranslation assay

WT or NGPS cells were seeded in 12-well plates. The next day cells were transfected with plasmid pRM hRluc-hFluc D357X, where Asp^357^ (GAC codon) was replaced by a UGA nonsense codon in the firefly luciferase (Fluc) transcript (*34*), using TransIT-2020 (Mirus #MIR5404) according to the manufacturer’s protocol. 30 hours post-transfection cells were lysed in 200 μL/well passive lysis buffer (Promega #E1941) for 15 minutes at r.t. under gently rocking. Lysates from each 12- well were collected in 3 wells of a black 96-well plate (PerkinElmer #6005182) on ice. The activities of firefly and sea pansy (*Renilla*) luciferases were measured sequentially using reagents of the dual-luciferase reporter (DLR) system (Promega #E1910) with a ClarioStar plate reader (BMG Labtech Ltd., Aylesbury, U.K.). In brief, firefly luciferase activity was assayed through addition of 60 μL/well Luciferase Assay Reagent II. Next, firefly luciferase activity was quenched and *Renilla* activity measured by addition of 60 μL/well *Renilla* substrate in Stop&Glo buffer. As a positive control for translation errors cells were treated for 24 hours with 0.5 mg/mL G418 (Gibco # 10131035) prior to assaying.

#### Statistics

Statistical analysis was done using Graphpad Prism v9. The statistical test used is indicated in the figure legends. Asterisks in the figures correspond to *p*-values as follows: * 0.01 ≤ *p* < 0.05, ** 0.001 ≤ *p* < 0.01, *** 0.0001 ≤ *p* < 0.001, **** *p* < 0.0001. ns indicates a non-significant statistical comparison.

#### Genome wide association studies (GWAS)

Target genes from the whole genome screen were integrated with common variant genome-wide association studies (GWAS) and associated functional annotations, pertaining to coding variants, expressions quantitative trait loci (eQTL) datasets and proximal enhancers.

For the common variant GWAS, we used data on body mass index (BMI, n= 806,834) and waist-hip ratio (WHR) adjusted for BMI (n=694,649) from the GIANT study (*51*), the GLGC triglycerides study (*52*) (n=1,253,275) and the GEFOS estimated bone mineral density (eBMD, n=426,824) study (*53*) and only variants with a minor allele frequency >0.1%. For each of the target genes and each phenotypic trait, genes were annotated based on proximity to genome-wide significant signals (p < 5×10^-8^), in 1Mb windows; 500kb up- or downstream of the genes start or end site. As most GWAS signals are intronic or intergenic, we overlayed these associations with other functional datasets to understand whether the associated variants can be causally linked to changes in the identified genes’ regulation or expression.

First, we calculated genomic windows of high linkage disequilibrium (LD; R^2^ > 0.8) for each given signal and mapped these to the locations of known enhancers for the target genes, using the activity-by-contact (ABC) enhancer maps (*32*), to indicate whether the genomic variants associating with the traits of interest directly changed the sequence of enhancers for the genes in question. We then performed colocalisation analyses between the four GWAS and expression quantitative trait loci (eQTL) data for genomic variants reaching at least a suggestive level of significance in the GWAS (*p* < 5×10^-5^), using the SMR and HEIDI tests (v1.02, (*54*)) and blood gene expression level data from the eQTLGen study (*55*). In doing so we essentially matched the pattern of association observed between the identified genomic variants and the GWAS outcome to the association towards the measured transcript level changes for the identified genes, to approximate the direct effect of the associated variants on the target gene expression. We considered gene expression of a gene to be influenced by the same genomic variation as that seen in the GWAS, if the FDR-corrected *p*-value for the SMR-test was p<0.05 and the *p*-value for the HEIDI test was >0.1%. Both of these sets of results allow us to draw methodological hypotheses about how genetic variants at the identified loci, can cause changes in the observed phenotypic traits, by directly affecting the genes regulation and expression patterns.

Finally, outwith transcriptional changes, we wanted to identify potential coding variants that might also associate with variation in the four phenotypes, indicating more direct protein-level consequences to the identified genes. To do this, we collapsed common coding variants within each of the identified genes and calculated gene-level associations towards each of the four traits, using a gene-level MAGMA analysis (*56*). Genes exhibiting an FDR-corrected MAGMA *p*-value <0.05 were considered significant.

#### *C. elegans* maintenance

BN808 *baf-1(bq19[G12T]) III* was generated by backcrossing *baf-1(G12T)* mutants (a kind gift from Dr Christian Riedel) six times with the N2 wild type strain. The *baf-1(bq19[G12T])* allele contains two nucleotide substitutions at position 34-35 relative to the start codon (GG→AC) introduced by CRISPR/Cas9 genome engineering. BN1336 *yc32[gfp::lmn-1] I; baf- 1(bq19[G12T]) III* was generated by crossing BN808 and UD484 (*57*) and balanced with hT2 [bli- 4(e937) let-?(q782) qIs48] (I;III)). BN1389 *knuSi221[fib-1p::fib-1::gfp + unc-119(+)] II; baf- 1(bq19[G12T]) III* was obtained by crossing BN808 and COP262 (*58*). Strains were maintained at 16°C on solid Nematode Growth Medium (NGM) plates seeded with *Escherichia coli* OP50 bacteria (*59*).

#### *C. elegans* nucleolar size measurement

Nucleolar area was quantified in the COP262 and BN1389 strains using the FIB-1::GFP reporter. Larvae were raised at 16°C and synchronized by picking L4s and further incubated for 24 hours at either 16 °C or 25°C. Animals were anaesthetized in a drop of 10 mM levamisole on top of soft agar pads as described (*60*). Stacks of confocal images were acquired using a Nikon Eclipse Ti microscope equipped with Plan Fluor 40×/1.3 and Plan Apo VC 60×/1.4 objectives and an A1R scanner using a pinhole of 1.2 airy units. Nucleolar areas were measured after creating binary masks with Fiji software from the original images (*61*).

#### *C. elegans* RNAi

For RNA interference, worms were fed with *E. coli* that express double-stranded RNA (dsRNA). Bacterial clones were obtained from a genome wide RNAi library (*62*) except clones corresponding to *mel-28*, *npp-6*, *npp-20*, *rpn-9* and *rpl-13* that were obtained from alternative sources (*63–65*). RNAi plasmids for *csk-1*, *daf-4*, *gft-2H3*, *src-1*, *ced-9* and *mrpl-36* were constructed as described (*66*) using the PCR primers listed in Table S2. PCR fragments were inserted into plasmid pL4440 *npp-15* (*64*) after digestion with XhoI (NEB R0146S) and SpeI (NEB R3133S), ligation and subsequent transformation into *E. coli* DH5α. Plasmids from ampicillin resistant colonies were confirmed by restriction digestion, transformed into *E. coli* HT115 and plated on LB with ampicillin and tetracycline.

For RNAi experiments, 12-15 homozygous *yc32[gfp::lmn-1]; baf-1(bq19[G12T])* animals were transferred at L4 stage to RNAi plates containing dsRNA-producing *E. coli* HT115, 1 mM IPTG and 100 µg/ml ampicillin and incubated at 20°C. After 18 hours, young adults were transferred to fresh RNAi plates (3 animals per plate; nine worms in total per replica) and incubated for 6 hours at 20°C. The adults were then removed and rescue of lethality was determined by counting the number of unhatched embryos and viable offspring after 24 hours at 20°C.

#### Statistical analysis of *C. elegans* data

Data were analyzed in R Studio (1.3.1093 (*67*); running R 4.0.2 (*68*)) and represented with base plotting. Mann-Whitney and *t*-tests were performed in R Studio and adjusted for multiple comparison (Benjamini & Hochberg method) when relevant.

---

**Fig. S1.**
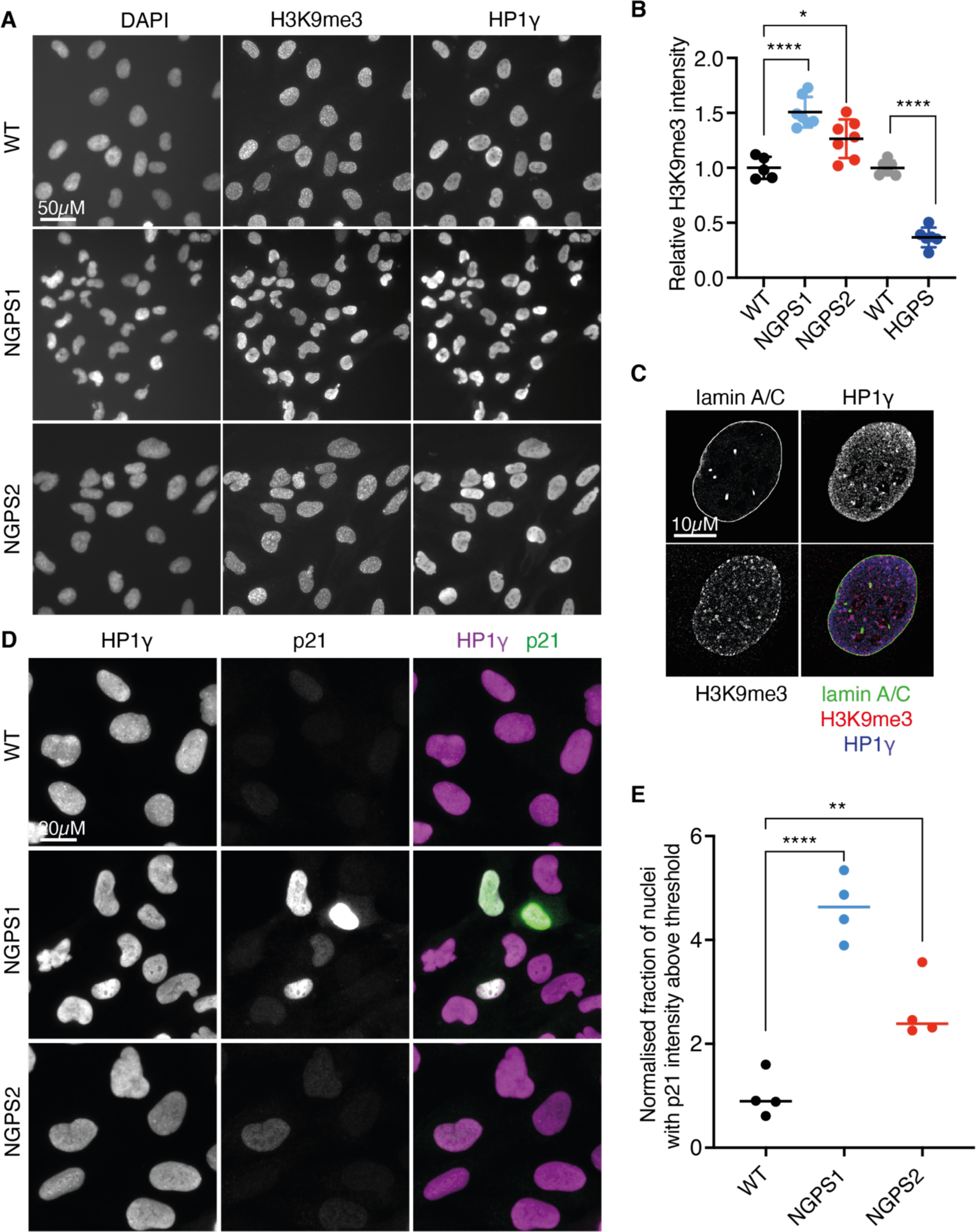
Heterochromatin marks and p21 are increased in NGPS fibroblasts. (**A**)Representative immunofluorescence images of WT, NGPS1 and NGPS2 fibroblasts stained for the heterochromatin marks H3K9me3 and HP1γ as indicated. DNA is stained with DAPI. Images were obtained with the high-content microscope. (**B**) Quantification of H3K9me3 intensity in the indicated progeria cell lines, compared to the corresponding WT. Data was obtained using HCS studio and represents the relative expression, in 2 experiments, each averaging 500 nuclei in 3 wells. Student’s *t*-tests were used for pairwise comparisons. (**C**) Representative super-resolution immunofluorescent staining of a WT cell nucleus showing that HP1γ overlaps with H3K9me3 and is therefore associated with heterochromatin domains in fibroblasts. (**D**) Representative wide-field immunofluorescence images of HP1γ and p21 in WT, NGPS1 and NGPS2 fibroblasts. (**E**) Quantification of the fraction of nuclei showing high level of p21 in the indicated cell lines, compared to the WT. Data was obtained using HCS studio and represents the relative fraction of nuclei with p21 intensity above a set threshold value, as obtained in 4 experiments, each measuring 500 nuclei in 3 wells. Student’s *t*-tests were used for pairwise comparisons.

**Fig. S2:**
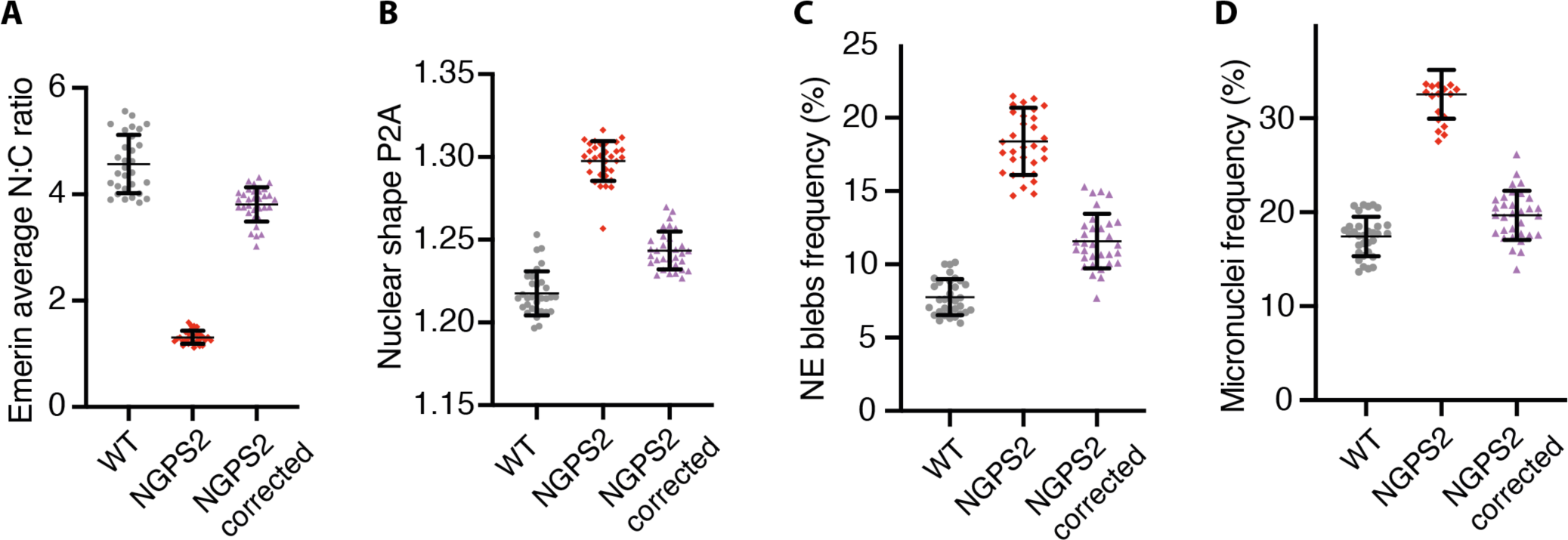
Validation of the screening phenotypes specificity using NGPS2 corrected isogenic cells. (**A-D**) Quantification of the indicated phenotype in WT, NGPS2 and NGPS2 corrected cells, in which the BAF A12T mutation has been reversed using CRISPR/Cas9 ^(14)^. P2A is a perimeter to area (P2A) analysis of nuclear shape. All the data were obtained using HCS Studio software. Each data point is the average value measured over 500 cells; lines indicate the average ± SD.

**Fig. S3.**
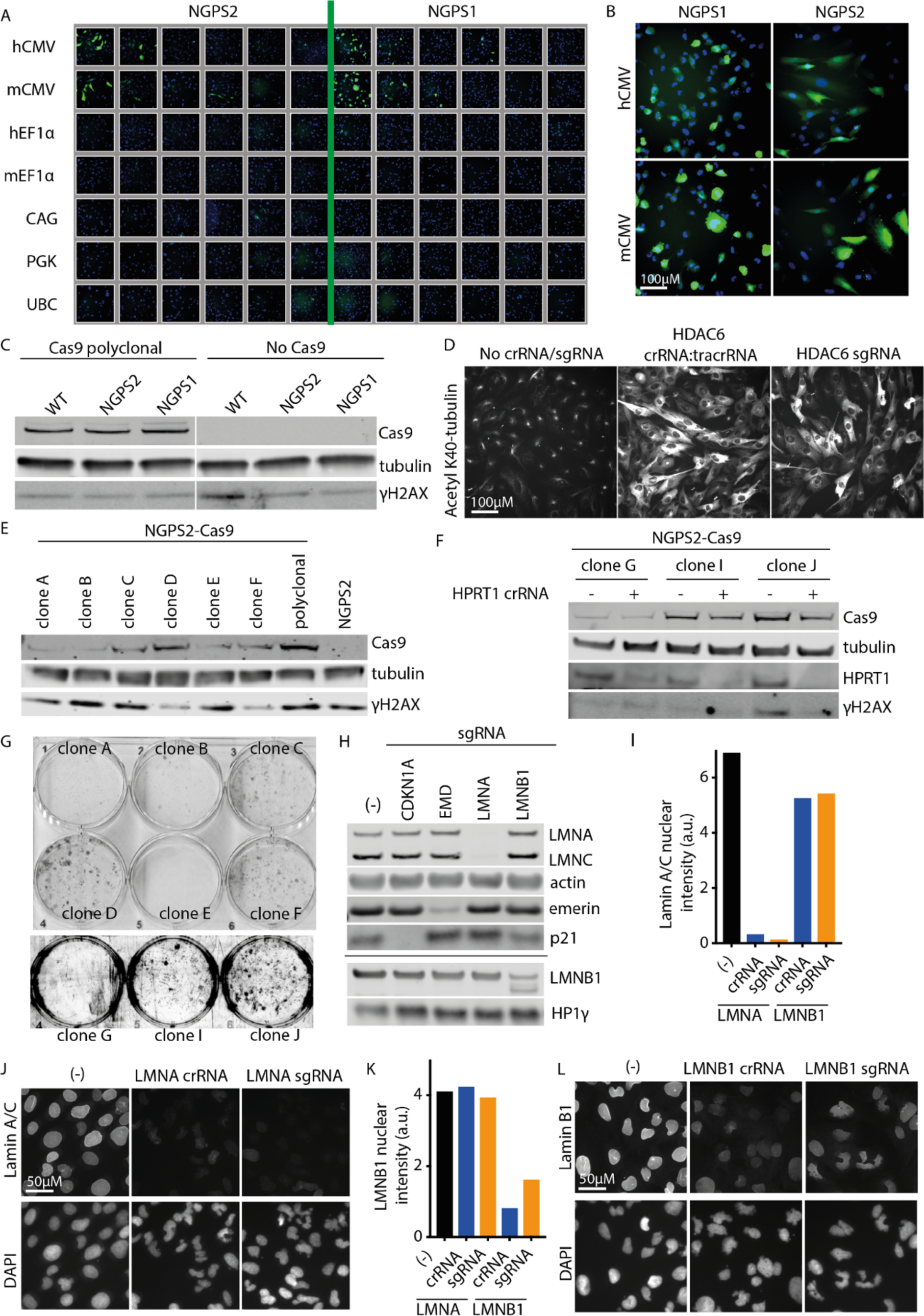
Engineering Cas9 expression and assessing its efficiency in NGPS fibroblasts. (**A**) A GFP construct was transduced in a SMARTchoice promoter selection plate in both NGPS1 and 2 cell lines to identify the promoter leading to the highest protein expression. Nuclei were counterstained with DAPI and the plate was imaged with the high-content microscope. (**B**) Zoom in from images obtained in (A) showing that GFP expression was highest in NGPS fibroblasts with the hCMV promoter. (**C**) Representative western blotting showing that Cas9 expression does not increase γH2AX levels in WT and NGPS polyclonal populations. (**D**) Representative immunofluorescence images showing the increase of acetyl-tubulin on lysine 40 (K40), used as a readout for HDAC6 depletion efficiency following cr:tracr or sgRNA transfection in polyclonal WT-Cas9 cells. The images illustrate efficient but heterogeneous knock-out using both reagents. (**E**) Representative western blot showing the expression level of Cas9 and of the DNA double strand break marker γH2AX in the parental NGPS2 cell line, in the NGPS2-Cas9 polyclonal population or in individual clones grown from single cells. (**F**) Representative western blotting showing the efficiency of HPRT depletion in the indicated NGPS2-Cas9 clones 72 hours post-transfection. (**G**) Resistance to 6-thioguanine was assessed in the indicated clones following HPRT depletion as shown in (F), to identify clones with the highest Cas9 cutting efficiency. (**H**) Representative western blotting showing the efficiency of the indicated sgRNA-mediated gene depletion in NGPS1-Cas9 cells 72 hours post-transfection. (**I-L**) Comparison of the knockout efficiency of the nuclear envelope proteins emerin, lamin A/C and lamin B1 (LMNB1) following transfection with a pool of 3 crRNAs or 3 sgRNAs in WT-Cas9 cells and quantified using high-content microscopy. Due to limited reagent availability the data represent a single experiment in which 1000 nuclei were imaged per condition.

**Fig. S4.**
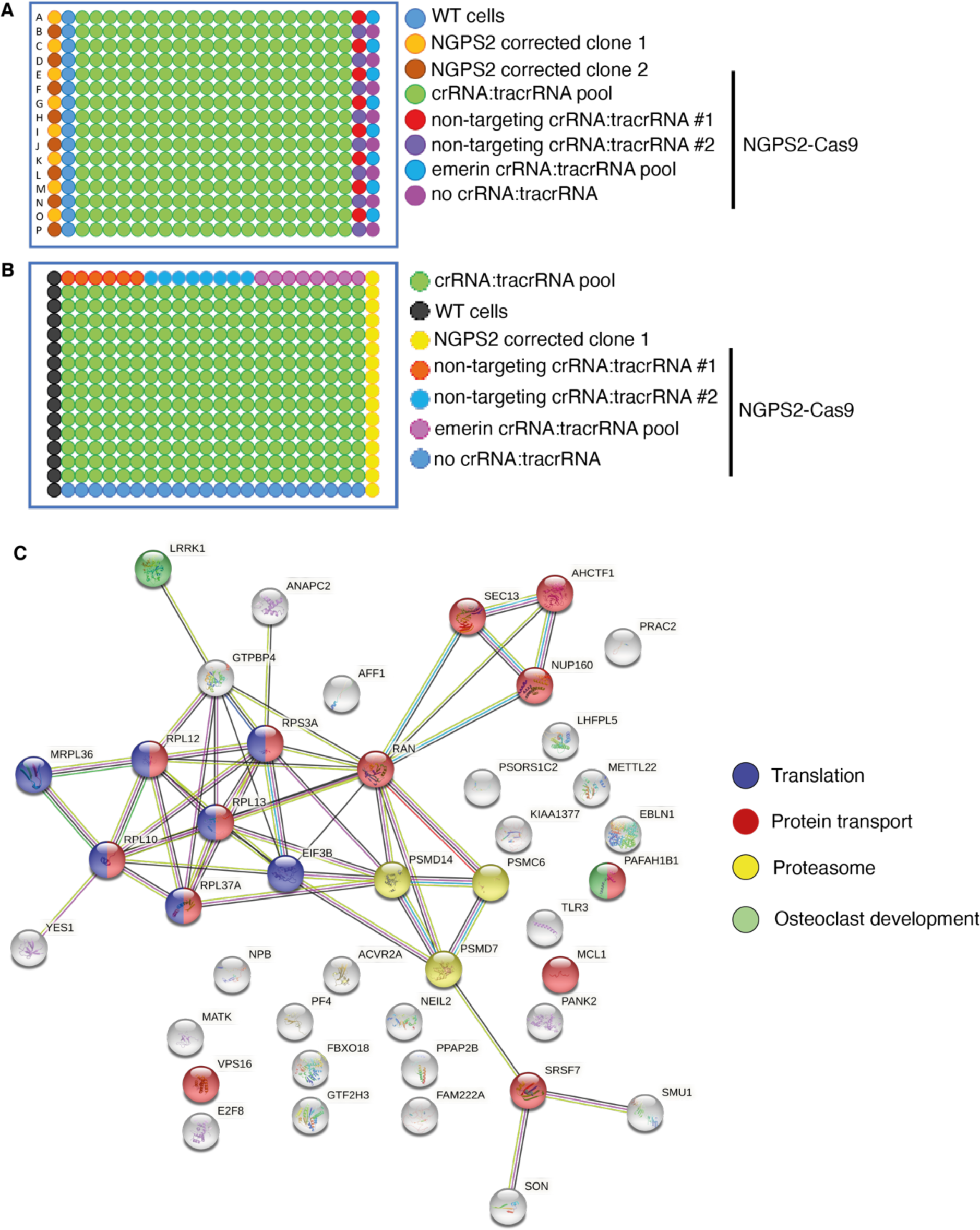
Screening plates layout and validation. (**A**) 384-well plate layout for the whole genome primary screen. WT cells as well as NGPS2 corrected cells (two separate clones) were used as control cells. Non-targeting crRNAs as well as wells without any crRNA were used as negative controls for the crRNA and transfection toxicity respectively. Emerin crRNA was used as a positive control for the transfection efficiency on each plate. (**B**) 384-well plate layout for the validation screen. Assay wells now contain a single crRNA:tracrRNA. Controls are as in (A). (**C**) STRING protein-protein interaction diagram. Gene ontology analysis revealed proteins involved in processes of translation (blue nodes), protein transport (red nodes), proteasome (yellow) and osteoclast development (green).

**Fig. S5.**
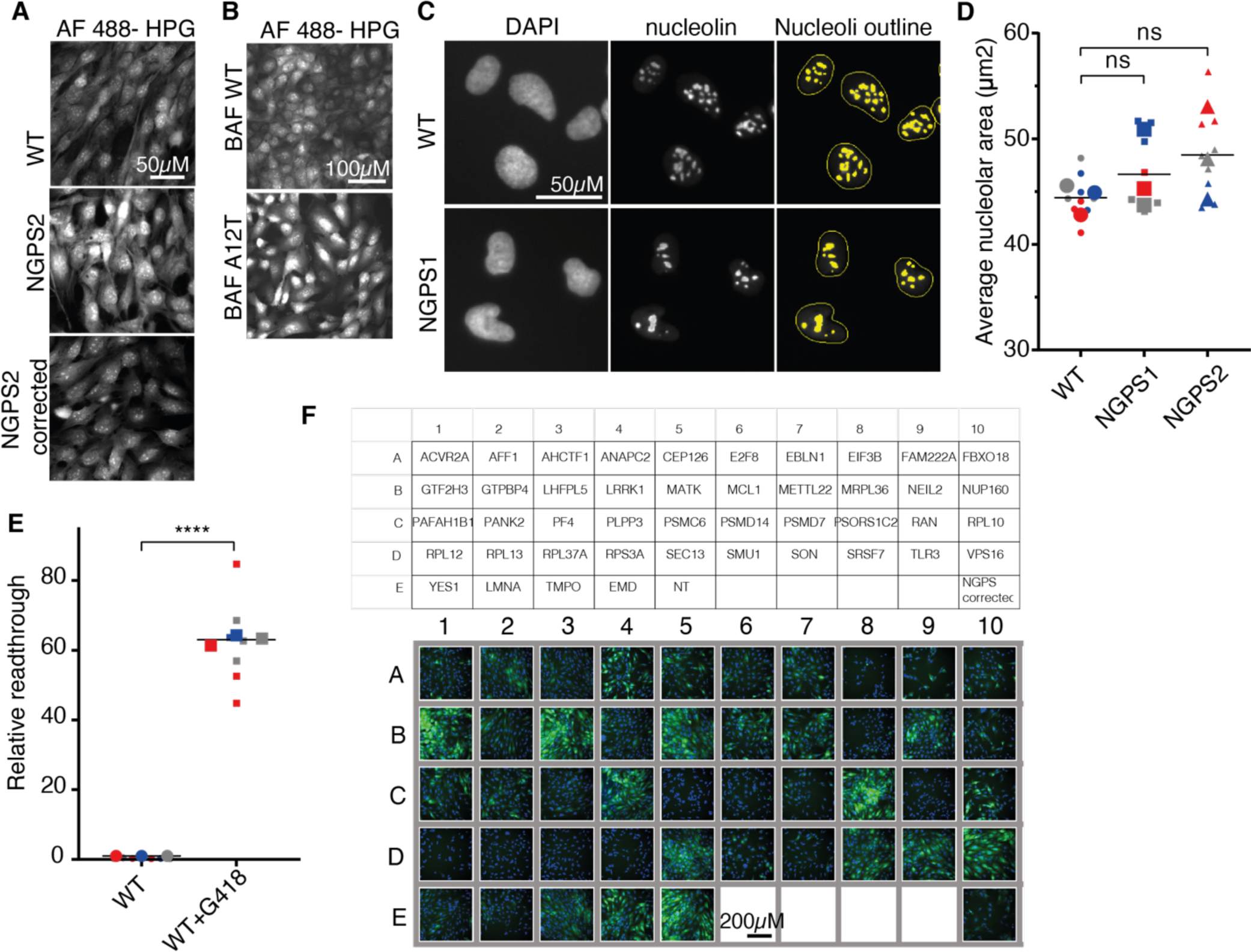
Enhanced protein synthesis in NGPS cells correlates with nuclear envelope abnormalities. (**A-B**) Nascent protein synthesis assay using HPG incorporation followed by labelling using a “click” reaction with Alexa Fluor 488 azide (AF 488). Representative immunofluorescence images of AF 488 HPG in WT, NGPS2 or NGPS2 corrected (mutation reversed by CRISPR/Cas9) (**A**) or in WT fibroblasts expressing a BAF WT or BAF A12T construct (**B**). (**C**) Immunofluorescence images of nucleolin in the indicated cell lines used to identify and outline the nucleoli (yellow) using the high-content microscope and analysis software. DAPI staining was used to define the nuclear mask. (**D**) Average nucleolar area quantified in NGPS1 and NGPS2 compared to WT fibroblasts by high-content microscopy. Separate colors indicate individual experiments. In each experiment 500 nuclei were analyzed in 3 separate wells with the larger symbols indicating the average measured value for each experiment. Results are compared using paired *t*-test. (**E**) Translation error rate measured as an increased read-through using a dual luciferase assay. G418 was used as a positive control for the induction of translational errors (*48*). Results are derived from the ratio hFluc/hRluc and given as fold induction. Three independent experiments are indicated in a superplot, with separate colors indicating individual experiments and larger symbols indicating the average for each experiment. Results are compared using an unpaired *t*-test. (**F**) Top: plate layout indicating the siRNA-targeted genes used in the nascent protein synthesis assay in NGPS2 cells. NT represents a non-targeting siRNA control. Well E10 was seeded with NGPS2 corrected cells as WT control. Bottom: Fluorescence intensity of HPG AF 488 imaged with the high-content microscope showing reduction of protein synthesis in NGPS2 cells upon depletion of most genes, compared to the non-targeting siRNA.

**Fig. S6.**
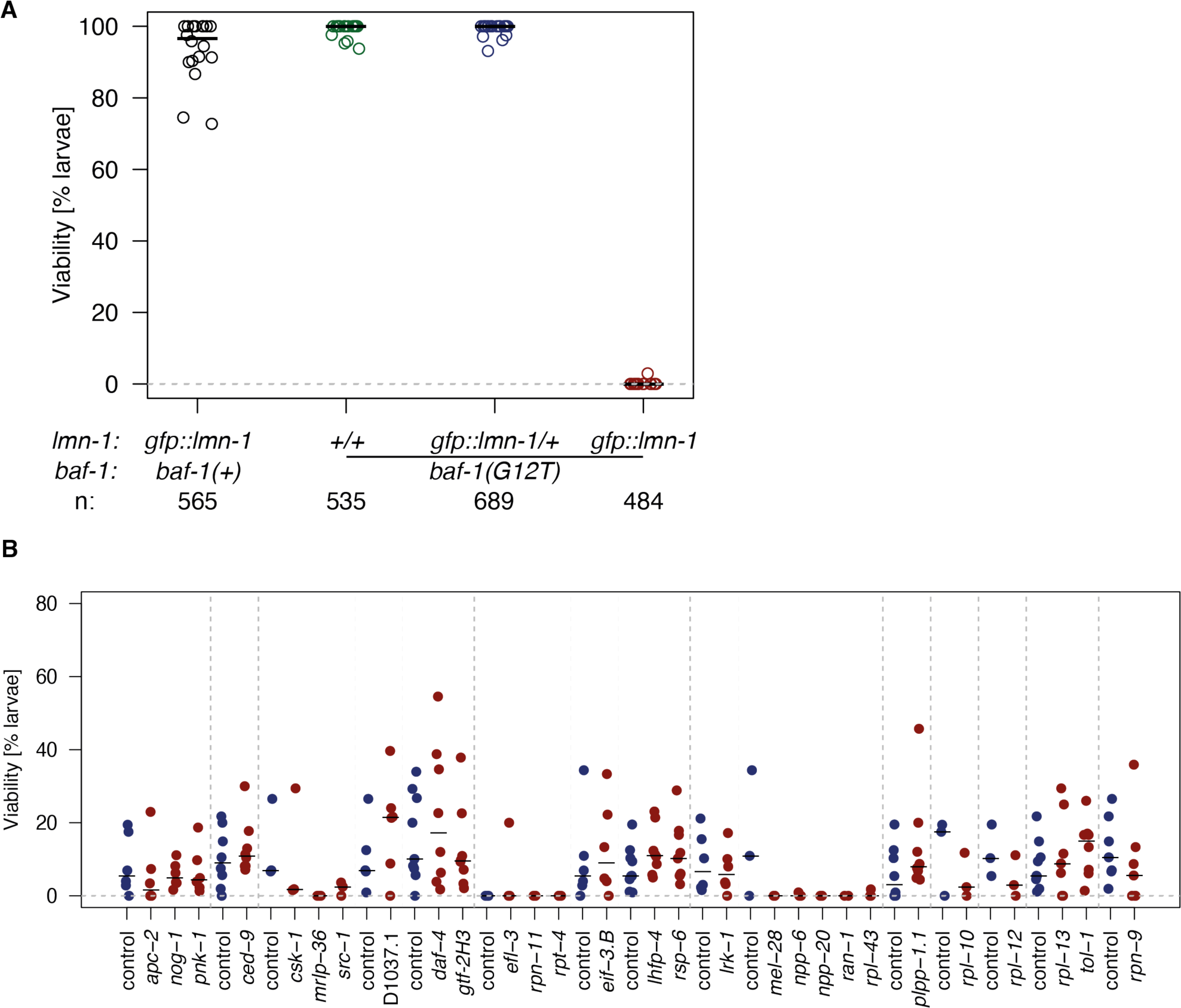
Evaluating the effect of candidate genes knock down in an NGPS *C. elegans* model. **(A)** Viability was determined for BAF-1 WT worms expressing endogenously tagged GFP::LMN- 1 (black circles) and for *baf-1(G12T)* worms carrying either no *gfp::lmn-1* allele (green circles), a single allele (blue circles) or the *gfp::lmn-1* alleles in homozygosity (red circles). Each circle corresponds to an individual plate and total number of eggs analyzed (n) is indicated. Note that this experiment was performed with *E. coli* OP50 as food source, which induces higher lethality in *gfp::lmn-1; baf-1(G12T)* animals as compared to *E. coli* HT115 used in RNAi experiments. **(B)** Candidate genes were knocked down by RNAi and tested for suppression of lethality in *gfp::lmn- 1; baf-1(G12T)* animals. Each point corresponds to the percentage of eggs developing into larvae from a single plate with 50-100 eggs; 2-13 plates were evaluated in 1-4 experiments for each gene. Black lines indicate medians. Vertical dashed lines separate each set of control and test plates; no significant suppression was observed in any of these cases (Mann-Whitney test).

**Table S1.** List of antibodies and reagents used in Western blotting and immunofluorescence.

| **Name** | **Manufacturer** | **Catalogue number** | **Application(s)** | **dilution** |
| --- | --- | --- | --- | --- |
| Anti-emerin | ProteinTech | 10351-1-AP | WB and IF | 1:1000 (WB)<br>1:750 (IF) |
| Anti-lamin A/C | Santa Cruz Biotechnology | sc-7292 | WB and IF | 1:1000 |
| Anti-β-actin | Cell Signalling | 3700 | WB | 1:4000 |
| Anti-LAP2 | BD Biosciences | 611000 | IF | 1:1000 |
| Anti-α-tubulin | Sigma | T9026 | WB | 1:3000 |
| Anti-lamin B1 | Santa Cruz Biotechnology | sc-365214 | WB and IF | 1:1000 |
| Acetyl-α-tubulin (K40) | Cell Signalling | 5335 | WB and IF | 1:1000 |
| Anti-phospho-histone H2A.X (S139) | Millipore | 05-636-I | WB | 1:200 |
| Anti-53BP1 | Bethyl | A300-272A-M | IF | 1:500 |
| Anti-p21 | Cell Signalling | 2947 | WB and IF | 1:1000 |
| Anti-SIRT7 | Santa Cruz Biotechnology | sc-365344 | WB | 1:200 |
| Anti-RAN | BD Biosciences | 610340 | IF | 1:1000 |
| Anti-HP1γ | Santa Cruz Biotechnology | sc-398562 | WB and IF | 1:2000 |
| Anti-H3K9me3 | Abcam | ab8898 | WB and IF | 1:2000 |
| Anti-H3K79me2 | Abcam | ab3594 | WB | 1:1000 |
| Anti-Cas9 | Biolegend | 698301 | WB | 1:1000 |
| Anti-HDAC6 | Abcam | ab133493 | WB | 1:1000 |
| Anti-HPRT1 | Genetex | GTX113466 | WB | 1:1000 |
| Anti-nucleolin | Genetex | GTX13541 | IF | 1:1000 |
| DAPI | EMP Biotech | F-0410-M0001.0-001 | IF | 1:5000 |
| Alexa Fluor 488 anti-mouse IgG<sub>2b</sub> | Thermo Fisher Scientific | A21141 | IF | 1:1000 |
| Alexa Fluor 568 anti-mouse IgG<sub>1</sub> | Thermo Fisher Scientific | A21124 | IF | 1:1000 |
| Alexa Fluor 647 anti-rabbit | Thermo Fisher Scientific | A21241 | IF | 1:1000 |
| Alexa Fluor 488 azide | Thermo Fisher Scientific | A10266 | IF (Click reaction) | 1:1000 |
| IRDye 680RD anti-rabbit | Licor Biosciences | 925-68073 | WB | 1:15000 |
| IRDye 800CW anti-mouse | Licor Biosciences | 925-32212 | WB | 1:12000 |
| Prolong Gold | Thermo Fisher Scientific | P10144 | IF | none |

**Table S2.**
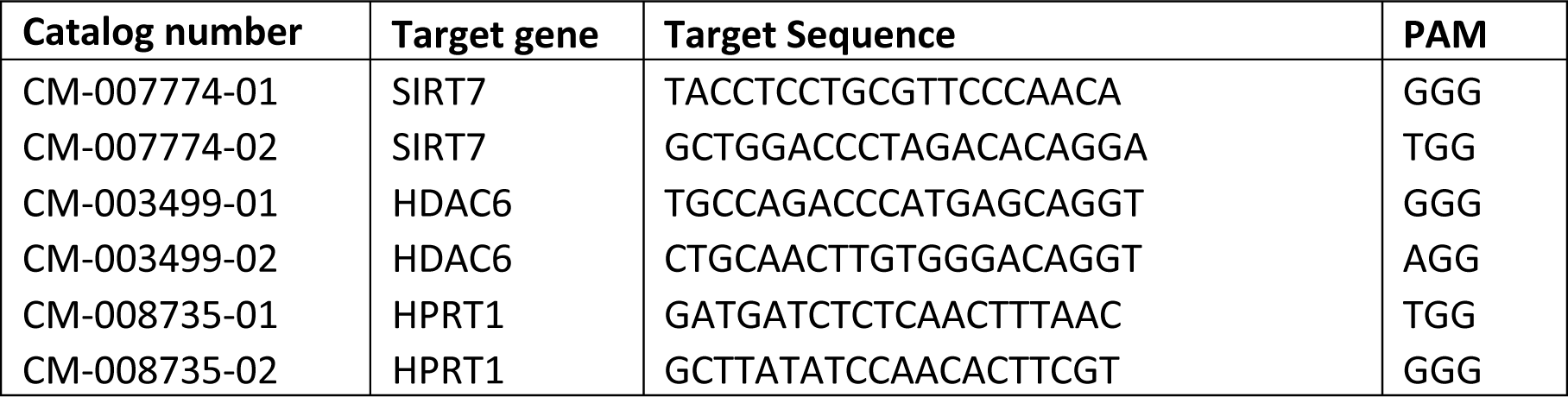
crRNA sequences used for CRISPR efficiency testing. (See Fig. 2J and Fig. S3D, F). All crRNAs were obtained from Horizon Discovery.

**Table S3.**
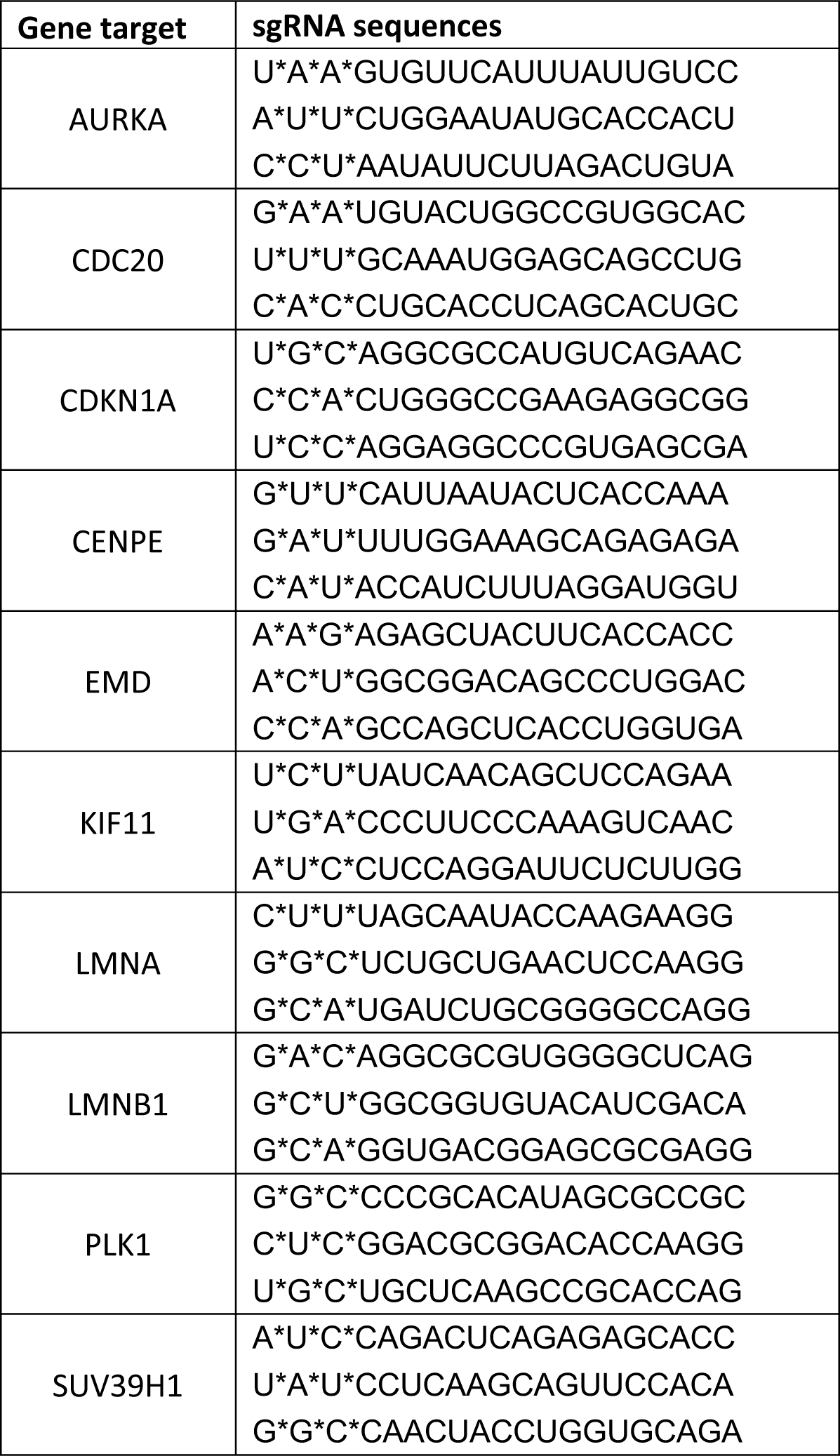
sgRNA sequences used in CRISPR efficiency testing experiments. Each gene was targeted by a pool of 3 sgRNA sequences. See Fig. 2K-L and Fig. S3H-L.

**Table S4.**
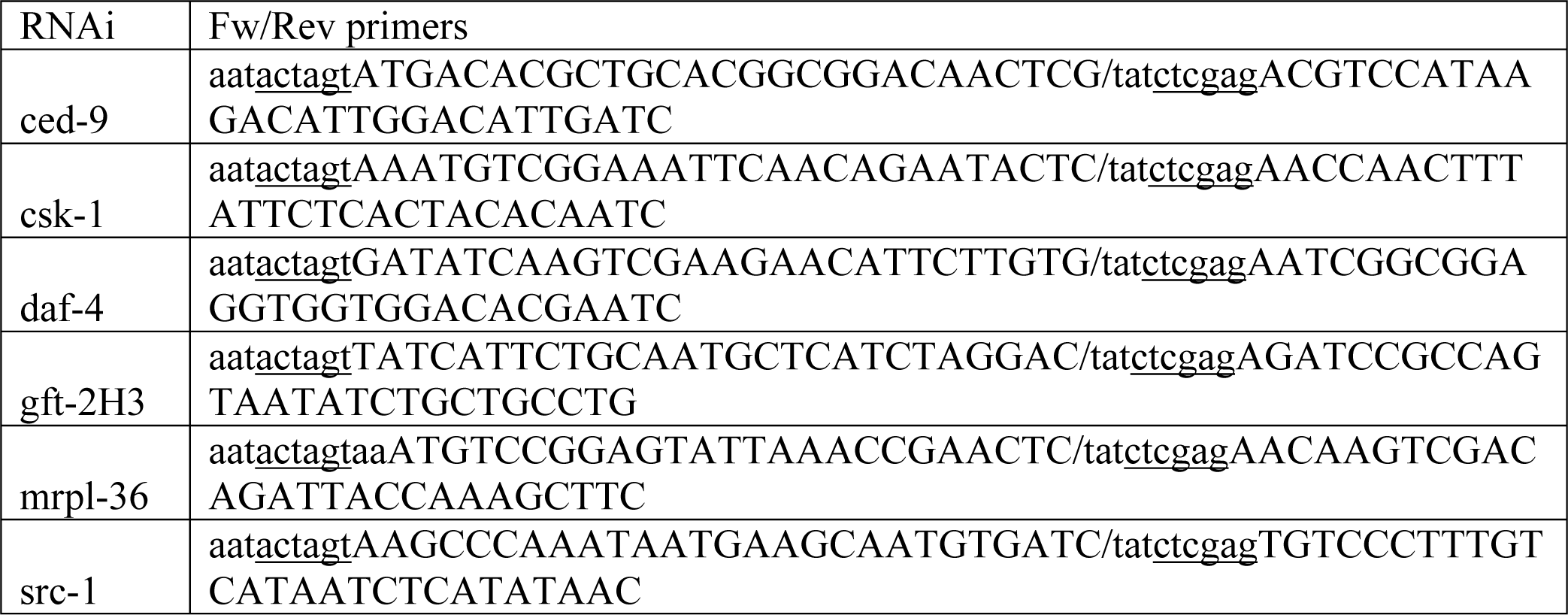
PCR primers used in *C. elegans* RNAi experiments.

## Data S1. (separate files)

Data S1 includes Tables S5-S12:

Table S5: Mini library crRNA sequences for CRISPR efficiency testing (Fig. 2L)

Table S6: Whole genome primary screen crRNA sequences

Table S7: Primary screen results

Table S8: Primary screen hits

Table S9: Validation screen crRNA sequences

Table S10: Validation screen results

Table S11: Mini library siRNA sequences for the nascent protein synthesis assay (Fig. 4G)

Table S12: Overlap of validated hit genes with human genetic and functional datasets (GWAS)

